# Coupling of electron-bifurcation modules powers aromatic ring reduction beyond the biological redox window

**DOI:** 10.64898/2026.04.10.717634

**Authors:** Lena Appel, Anuj Kumar, Davide Tamborrini, Tomas C. Pascoa, Kanwal Kayastha, Ulrich Ermler, Tim Kettler, Stefan Bohn, Tristan Reif-Trauttmansdorff, Benjamin D. Engel, Jan M. Schuller, Matthias Boll

## Abstract

Microbial degradation of ubiquitous aromatic compounds is central to the global carbon cycle and bioremediation, yet the intrinsic stability of aromatic rings poses a major barrier to their breakdown. Under strictly anaerobic conditions, class II benzoyl-CoA reductases (BCRII) catalyse the key step of this process, a Birch-like reduction of the aromatic ring to a cyclic diene at a tungsten cofactor. This reaction operates beyond the redox limits of conventional biological electron transfer, yet the mechanism by which BCRII generates such extreme reducing power has remained unclear. Here, high-resolution cryo-electron microscopy, *in situ* cryo-electron tomography, and enzymatic analyses reveal that the one-MDa BCRII complex from *Geobacter metallireducens* links two distinct flavin-based electron-bifurcation modules, previously characterised in hydrogenases and heterodisulfide reductases, to drive aromatic ring reduction. Reduced ferredoxin and NADH deliver electrons through sequential confurcation and bifurcation to the catalytic site, while cryo-electron tomography of native cells identifies electron-transferring flavoprotein as the second acceptor directing high-potential electrons into the respiratory chain. These results show that modular FBEB units can be hierarchically assembled to extend metabolic redox capacity, highlighting their versatility as adaptable components for electron transfer pathways.

## Introduction

The redox span of most biological electron-transfer reactions lies within a window of ∼1.3 V, bounded by the O_2_/H_2_O couple (E°′ ≈ +0.8 V) and ferredoxin (Fd _ox/red_; E°′ ≈ –0.5 V). Fds are small iron-sulfur (Fe–S) proteins that function as versatile redox carriers and are widely regarded as ancestral electron-transfer mediators^1^. Only few biological processes operate at potentials more negative than Fd, such as nitrogen fixation by nitrogenases^2^, activation of methyl-coenzyme M reductases for methane production^3^, or aromatic ring reduction by benzoyl-coenzyme A (CoA) reductases (BCRs)^4^. BCRs catalyse the dearomatisation of benzoyl-CoA to cyclo-hexa-1,5-diene-1-carbonyl-CoA (1,5-dienoyl-CoA), a key step in the anoxic degradation of aromatic compounds, which, after carbohydrates, represent the second most abundant class of biomolecules on Earth, primarily derived from lignin and fossil oils^5^. In anoxic environments, BCRs play a pivotal role in global carbon cycling and in the breakdown of homocyclic aromatic pollutants such as benzene, toluene, ethylbenzene, and xylene (BTEX) hydrocarbons^4,6^. Monitoring BCR-associated genes provides a powerful approach to evaluate microbial biodegradation potential in BTEX-contaminated anoxic ecosystems, as demonstrated during the Deepwater Horizon oil spill^6,7^.

Class I BCRs overcome the negative redox potential limit of around –0.5 V by coupling electron transfer to stoichiometric ATP hydrolysis, a strategy feasible for facultative anaerobes with a considerably high ATP yield^8,9^. In contrast, strictly anaerobic bacteria prevalent in marine or freshwater sediments utilise class II BCRs (BCRII), which are proposed to use flavin-based electron bifurcation (FBEB) to access low-potential chemistry at reduced energetic cost^10,11^. FBEB, together with the reverse electron confurcation, constitutes a mode of energy coupling, in which a mid-potential donor reduces a flavin cofactor, from which electrons bifurcate to a high-potential (exergonic) and a low-potential (endergonic) acceptor. Four unrelated FBEB enzyme families are known, and nearly all family members function to reduce Fd^12–14^. They play key roles in many globally relevant microbial processes such as methanogenesis, acetogenesis, anaerobic respirations, and fermentations.

BCRII is distinguished among FBEB enzymes by its proposed use of Fd_red_ as a mid-potential donor to drive the reductive dearomatisation of benzoyl-CoA at E°′ ≈ –620 mV^15,16^. The ∼1-MDa Bam[(BC)_2_DEFGHI]_2_ BCRII complex, characterised from the strictly anaerobic deltaproteobacteria *Geobacter metallireducens GS-15*^*10*^ and *Desulfosarcina cetonica*^*17*^, contains more than 50 redox cofactors and ranks among the most intricate metalloenzymes described. The complex is predicted to be organised into three functional modules: (i) the BamB subunit, part of the BamBC module, binds the active-site aqua-tungsten-bis-metallopterin (W-MPT-OH_2_) cofactor and is a member of the aldehyde:Fd oxidoreductase (AOR) family of W-MPT cofactor-containing enzymes. At the W-MPT-OH_2_ centre, aromatic ring reduction is initiated via hydrogen-atom transfer from the aqua ligand, proceeding through a Birch reduction-like radical mechanism^15,18^; (ii) the BamDEF module, related to FBEB HdrABC heterodisulfide reductase-MvhAGD hydrogenase components with BamE predicted to bind the bifurcating flavin cofactor^11,19^; (iii) the BamGHI module, related to NADH/Fd_red_-oxidising HydABC modules of electron bifurcating/confurcating hydrogenases, containing a second electron-bifurcating flavin cofactor^20–22^ **(Extended Data Fig. 1)**. The coexistence of two distinct FBEB modules within a single oxidoreductase supercomplex has not been previously reported, and their function in BCRII remains unclear. Despite this predicted modular architecture and the proposed FBEB-driven mechanism, electron bifurcation-mediated aromatic ring reduction has not been experimentally observed, likely owing to a suggested membrane association and the involvement of menaquinone (MK) as high-potential acceptor via as-yet unidentified redox components^10^. Accordingly, without structural insights, the mechanism by which BCRII harnesses reducing power to drive a Birch-like reduction remains elusive. Combining enzymatic assays with high-resolution cryo-electron microscopy (cryo-EM) and in-cell cryo-electron tomography (cryo-ET), we reveal how BCRII channels electrons through two distinct energy-coupling pathways to power aromatic ring reduction outside the conventional biological redox window.

**Fig. 1.**
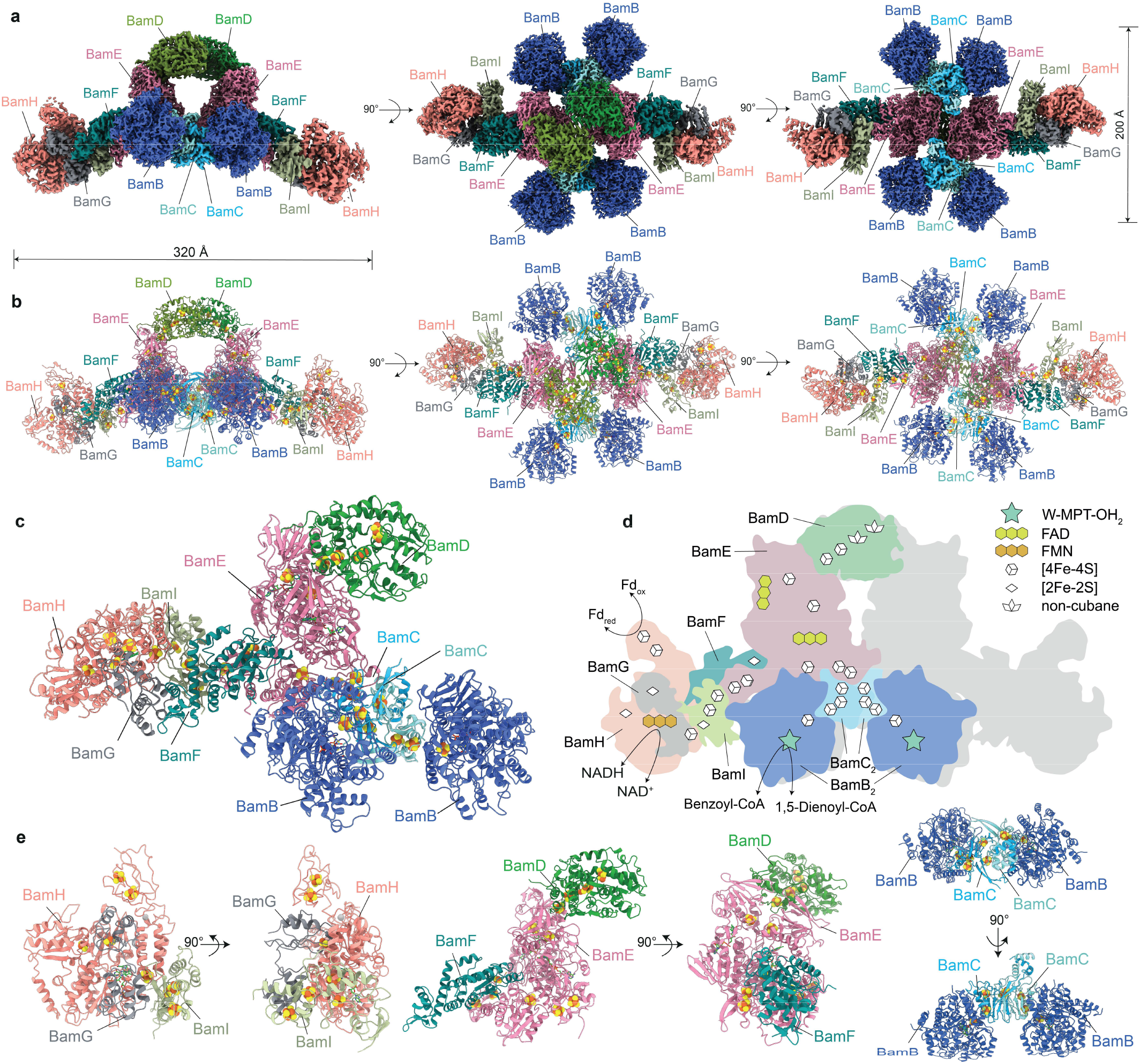
Molecular architecture of the BCRII complex. **a**, Segmented cryo-EM map of the dimeric BCRII complex, with subunits in distinct colours. **b**, Corresponding ribbon representation of the atomic model; Fe–S clusters shown as orange and yellow spheres. **c**, Atomic model of a functional BCRII protomer, with one copy each of BamG, BamH, BamI, BamF, BamE, BamD, and two copies of BamB and BamC. **d**, Schematic of the protomer showing subunit organisation and associated cofactors. **e**, Two views highlighting the three major modules: HydBC-like BamGHI, HdrABC-like Bam-FED, and AOR-type BamBC catalytic unit.

## Results and Discussion

### High-resolution cryo-EM structure of the BCRII complex

To elucidate the structure and mechanism of BCRII, we purified the native complex from *G. metallireducens* under strictly anoxic conditions^7,14^. BCRII preparations were homogeneous, exhibited the predicted subunit composition and cofactor complement, and retained 1,5-dienoyl-CoA-(reverse reaction) and NADH-dependent oxidoreductase activities, consistent with intact, functional complexes **(Extended Data Fig. 2a,c-e)**.

To define the molecular architecture of BCRII, we performed cryo-EM single-particle analysis. Owing to the complex’s oxygen-sensitivity^23^, samples were vitrified under strictly anaerobic, redox-controlled conditions. Prior to freezing, the complex was incubated with FMN and with the reductants NADH and Fd_red_, reduced by α-ketoglutarate:Fd oxidoreductase (KGOR)^10^, to establish physiologically relevant conditions and to capture potential redox-driven conformations. Reference-free 2D classification revealed that the BCRII complex adopts an Ω-shaped, dimeric architecture with C2 symmetry, featuring a rigid central core flanked by flexible peripheral arms at sub-stoichiometric levels. To obtain a structure encompassing both peripheral arms, a large dataset was collected to enable reconstruction of the fully assembled complex. Iterative 3D classification and focused refinement yielded a 1.9 Å map for the core, while the two flexible peripheral arms were resolved at 2.4 Å and 3.5 Å, respectively **(Extended Data Fig. 3-5; Extended Data Table 1)**. The locally refined maps were combined into a composite holo-complex map, allowing construction of a full atomic model of the BCRII assembly **(Fig. 1a-e)**. Each arm comprises a BamGHI assembly, whereas the core of the BCRII complex is formed by the BamED subunits. BamF, an MvhD-homologous adaptor, links the BamGHI and BamED assemblies^24^. At the periphery of the core, four BamBC catalytic modules surround the BamDE subunits forming the distal catalytic units of BCRII. The complete occupancy of the flavins, Fe–S clusters, and W-MPT cofactors was in line with biochemical analyses, indicating a fully functional BCRII complex.

### BamGH is the confurcating electron input module

Consistent with homology analyses, BamGH adopts the canonical architecture of the HydBC-dimer of FBEB hydrogenases catalysing the reversible reduction of NAD^+^ and Fd by H_2_. It is thus designated as an electron confurcation unit, accepting electrons from Fd_red_ and NADH. BamH forms the central scaffold of the module, displaying the conserved signature features of a HydB-bifurcating subunit^13,22,25,26^ **(Fig. 2a**,**b; Extended Data Fig. 6)**. Its architecture comprises an N-terminal thioredoxin (Trx)-domain containing the H3 [2Fe–2S] cluster, a core domain that binds the electron-confurcating FMN and NAD(H), and a C-terminal Fd-like domain harbouring the H1 and H2 [4Fe–4S] clusters, which serve as acceptors for external Fd_red_. The C-terminal domain is tethered to the core via a flexible linker, structurally stabilised by a coordinated Zn^2+^ ion. The BamH core adopts a modified Rossmann fold with four β-sheets that accommodate FMN and NAD(H), both observed at full occupancy in the cryo-EM map **(Fig. 2a)**. The nicotinamide ring of NAD(H) forms a π-stacking interaction with the isoalloxazine ring of FMN, with an interplanar distance of 3.85 Å between the NADH-C4 and the FMN-N5, suitable for direct hydride transfer. Adjacent to the core, a four-helical bundle domain contains the H4 [4Fe–4S] cluster, from which electrons are transferred to the BamI clusters I1-I3.

**Fig. 2.**
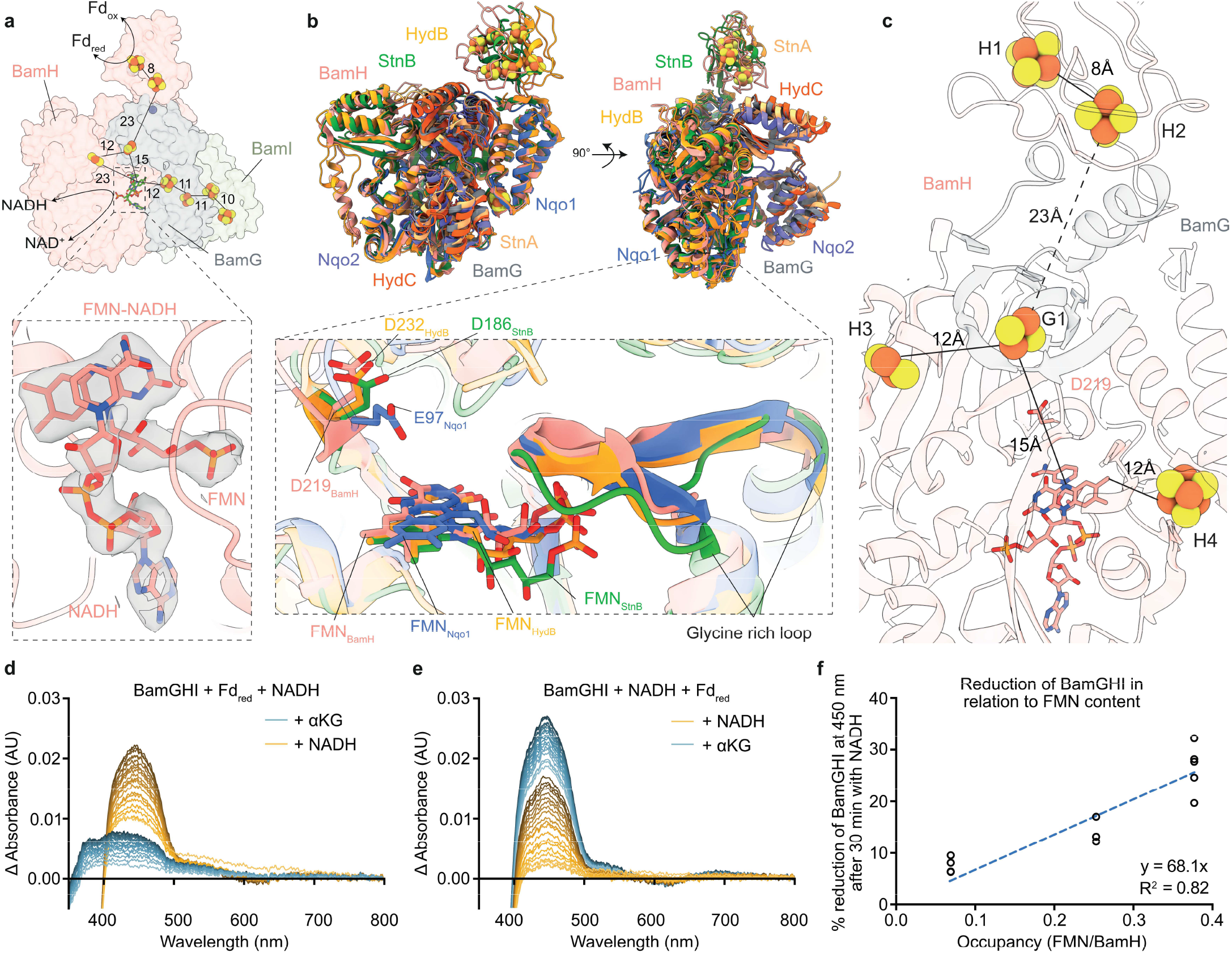
The BamGHI module performs a HydBC-type electron confurcation reaction. **a**, Surface representation of the BamGHI module, and binding of Fe–S clusters, FMN and NADH. FMN–NADH pair shown enlarged with cryo-EM density. **b**, Structural superposition of BamGH with HydBC-type complexes: HydBC (PDB 8A6T, RMSD 1.08 Å), StnAB (PDB 8OH9, RMSD 1.03 Å), and Nqo1/2 (PDB 3IAS, RMSD 1.05 Å). BamH contains a conserved FMN pocket stabilised by a glycine-rich loop; FMN is coordinated by Asp219 to stabilise a flavosemiquinone intermediate. **c**, Cartoon of BamGH showing cofactor distances; the G1–H2 separation exceeds typical electron-transfer range, proposed to be bridged by BamG C-terminal domain motion, repositioning G1 for directional electron flow towards FMN. **d**,**e**, UV/vis spectra showing time-dependent reduction of BamGHI by Fd_red_ (0.4 µM, reduced via αKG, KGOR, CoA) and NADH (100 µM), with reductants added in different orders. **f**, BamGHI reduction after 30 min plotted versus FMN occupancy. All distances in Å.

BamG, similar to HydC, associates with the module through contacts with BamH and with its N-terminal four-helical bundle engaging BamI. The G1 [2Fe–2S] cluster is embedded within the C-terminal Trx-domain **(Extended Data Fig. 6)**, positioning it in close proximity to the FMN-NADH binding site and the adjacent H2 and H3 cluster of BamH. These inter-subunit interactions create a contiguous network of cofactors, optimally arranged for efficient electron transfer from NADH and Fd_red_ to the downstream BamI Fe–S clusters.

In canonical EtfAB- and NfnAB-type FBEB systems, the bifurcating FAD is readily reduced to the hydroquinone (HQ) state, whereas one-electron reduction to the anionic semiquinone (SQ) is energetically disfavoured owing to its intrinsic instability^12,14,27^. By contrast, HydBC-type FBEB enzymes operate through a distinct mechanism in which the SQ state of FMN is stabilised^22,28^. Here, electron transfer from H_2_ to the acceptors NAD(P)^+^/Fd_ox_ is controlled by a redox-state dependent modulation of NAD(P)^+^ binding affinity and conformational gating between closed and open states. Although the details of energetic coupling remain unresolved, this mechanism enables alternating electron flow either to Fd_ox_ (endergonic) or NAD^+^ (exergonic), while preventing electron backflow. A flexible linker facilitates this process by dynamically coupling the Fd domain of HydB to the electron transfer pathway^22^.

In BamH, the key structural elements underlying the HydB-type FBEB mechanism are conserved. These include the flexible linker segment and a conserved aspartate residue (Asp219) positioned near the N5 of FMN, consistent with a role in stabilising the SQ state^22,26^ **(Fig. 2b)**. BamGH adopts the closed conformation, with FMN positioned ∼15 Å and ∼12 Å from the adjacent G1 and H4 clusters, respectively. In this state, the G1 cluster lies ∼23 Å from the H2 cluster, a separation too large to permit biological electron transfer **(Fig. 2c)**. Analogous to previous proposals for HydC^13,22,25^, we propose that conformational dynamics of the Trx-domain of BamG mediate transitions between closed and open states, with the open conformation positioning the H1 and H2 clusters in proximity to the G1 cluster.

To investigate BamGH-mediated flavin-based electron confurcation, with NADH and Fd_red_ serving as joint electron donors, we monitored Fe–S cluster reduction by NADH, Fd_red_, or both using UV/vis spectroscopy. For this purpose, the BamGHI subcomplex was heterologously produced in *Escherichia coli* under anaerobic conditions and purified to homogeneity, with an iron content of 25.9 ± 7.2 mol mol^−1^, consistent with five [4Fe–4S] and three [2Fe–2S] clusters **(Fig. 2a; Extended Data Fig. 2b-d)**. FMN occupancy was substoichiometric, as observed in other FBEB systems^12,29^; cluster reduction was therefore normalised to FMN content across biological replicates and benchmarked against complete reduction by dithionite **(Fig. 2f)**. Fd_red_ reduced only 20 ± 6 % of BamGHI **(Fig. 2d)**, which is consistent with the proposed electron-confurcation mechanism that blocks electron transfer from Fd_red_ via the H1 and H2 clusters to the G1 cluster in the absence of NADH. In contrast, NADH reduced 68 ± 4 % of BamGHI, whereas the combined reduction by NADH and Fd_red_ reached 84 ± 15 % **(Fig. 2e)**, representing an approximately additive effect of the two electron donors.

By analogy to the reverse HydBC-type electron bifurcation^22,30^, we propose a confurcation mechanism in which the H1/H2 clusters are readily reduced in the closed BamG-in conformation **(Extended Data Fig. 7)**. NADH binding induces a transition from the closed to the open BamG-out state, enabling electron transfer from the Fd_red_-reduced H1 and H2 clusters to the G1 and H3 clusters. Subsequent structural rearrangement to the closed state directs electron flow from NADH and the reduced G1 and H3 clusters towards the FMN and H4 cluster, thereby enforcing unidirectionality in this reverse confurcation mechanism. This mechanism allows NADH and Fd_red_ to act alternately as electron donors, sustaining efficient electron flow through the complex. Although the redox potential of NADH appeared sufficient to reduce the BamGHI clusters, Fd_red_ may become critical as joint electron donor at a downstream step, for example during the transition from one-to two-electron chemistry in the redox cofactors of the BamFE subunits.

### BamE bifurcates electrons to the BamD and BamB output modules

The BamGH confurcating unit is electronically coupled to the HdrA-type FBEB module BamE via BamI and the BamF adaptor **(Fig. 3a)**^19,24^. BamE retains the HdrA-type modular architecture, albeit with extensive domain permutation. Remarkably, BamE is composed of a fused HdrA dimer with each HdrA-like unit comprising a Trx reductase (TrxR)-like core domain complemented by N- and C-terminal regions and an inserted Fd domain (**Extended Data Fig. 6**). Both TrxR domains bind an FAD adjacent to the E4 and E7 [4Fe– 4S] clusters, respectively. The FAD in the N-terminal TrxR1 domain displays hallmark features of Hdr-type electron bifurcating flavins (B-FAD), including a conserved lysine (Lys280) positioned 3.4 Å from the N5 atom of the isoalloxazine ring, consistent with HQ stabilisation^19^. By contrast, the second FAD lacks key FBEB features and is not positioned at an electron-branching point, arguing against a role in electron bifurcation (NB-FAD) and instead supporting a primarily structural, rather than catalytic function of the TrxR2 domain **(Fig. 3b)**.

Despite the substantial structural adaptations, the overall electron-transfer wiring in BamE mirrors that of HdrA. Electrons are transferred from the I3 cluster to the E1 and E2 [4Fe-4S] clusters in the C-terminal Fd-like domain, while cluster E4 mediates transfer between the bifurcating FAD and the D1 cluster in BamD. Additional [4Fe-4S] clusters (E3, E5, and E6) connect the B-FAD to the Fe–S network of the BamBC active site.

The BamF adaptor – over 100 amino acids longer than previously described MvhD adaptors in HdrA-containing systems – comprises two Rossmann-fold domains and a winged helix–turn–helix motif that interfaces with BamI, forming a unique electron relay between the BamGH module and the BamED region. The BamF F1 [2Fe-2S] cluster accepts single electrons from the E2 cluster and relays them to B-FAD across a ∼28 Å gap, a distance incompatible with biological electron transfer.

Through iterative 3D classification and focused refinement of the BamGHIF region, we resolved two conformational states of BCRII, distinguished by a ∼28° rotation of the BamFGHI subcomplex and associated BamE regions **(Fig. 3c; Extended Data Table 1)**. Although differing in amplitude and direction from motions observed in methanogenic Hdr complexes, the resulting rearrangements of redox cofactors are mechanistically analogous^31^. Based on studies with HdrA complexes, state 1 (2.3 Å resolution) likely represents a pre-bifurcation configuration, in which the F1 cluster is oriented towards B-FAD at the ∼28 Å distance **(Fig. 3d; Extended Data Fig. 3, 8)**. State 2 (2.5 Å resolution) is defined by formation of a Cys59–Cys62 disulfide, which has been proposed to represent a post-bifurcation state in Hdr complexes^31^.

Electron transfer from the F1 cluster to B-FAD remains unresolved. By analogy to HdrA-type systems, thiol/selenol chemistry may facilitate this step. In methanogenic MvhD, a cysteine near the [2Fe–2S] cluster forms a regulatory disulfide enabling two-electron reduction via sequential one-electron steps^19^, reminiscent of Fd-dependent Trx reductases^32^. In BamF, this residue is a selenocysteine (Sec19), suggesting a possible selenenylsulfide intermediate. The F1 cluster may thus mediate the transition from one-to two-electron chemistry before transfer to B-FAD, potentially via the Cys59–Cys62 disulfide and a B-FAD–thiolate adduct. However, observed distances preclude direct thiol/selenol-disulfide exchange reactions **(Fig. 3d)**, and direct one-electron reduction of disulfides is energetically unfavorable with electrons derived from Fd_red_/NADH^33^. In summary, elucidating precise mechanistic details of electron transfer in HdrA-type FBEB systems represents an important avenue for future investigations.

Two-electron reduction of B-FAD triggers electron bifurcation, routing the high-potential first electron from the stabilised HQ to cluster E4 (towards BamD) and the low-potential second electron from the low-potential SQ intermediate to cluster E3 (towards BamB). E3 then channels low-potential electrons to the BamB active site, yet its ∼17 Å distance to B-FAD precludes direct transfer. Instead, the C59–C62 disulfide is ideally positioned, 3.4 Å from cluster E3 and 13.3. Å from B-FAD, for a linear single electron transfer from the B-FAD through the disulfide to cluster E3. Notably, the redox-potential of the Q/SQ redox couple in electron-bifurcating flavins is estimated to be –0.9 V or even lower^34^, and sufficiently low for single electron transfer via a disulfide to cluster E3. Transition to state 1 shortens the E3-E5 distance from 23 Å to 12 Å, thereby promoting low-potential electron transfer to the E5 and E6 clusters in the inserted Fd domain. This Fd domain connects to the low-potential output module Bam(BC)_2_, analogous to HdrA’s Fd domain, which reduces external Fd. The low-potential electrons are transferred through the three [4Fe-4S] clusters of BamC to the adjacent BamB subunits coordinating the ring-dearomatising active site W-MPT-OH_2_ cofactor and a [4Fe-4S] cluster^15^.

The high-potential electron is transferred to the E4 cluster in the TrxR domain, which associates with BamD, a subunit resembling a fusion of heterodisulfide reductase HdrBC^19^ **(Extended Data Fig. 6, 9a-c)** or HdrD^35^ subunits. In methanogens, electrons pass through two cubane [4Fe-4S] clusters in HdrC to reach two non-cubane [4Fe-4S] clusters in HdrB, reducing CoM-S-S-CoB^19^. In BamD, the CoM-S-S-CoB binding site is occupied by the His230, Trp325 and Tyr356, arguing against heterodisulfide binding. The two cubanes (D1, D2) and the proximal non-cubane cluster (D3) are conserved, whereas the distal non-cubane (D4) Fe−S cluster is distinct **(Fig. 3e)**. In *G. metallireducens* BamD, a cysteine-to-serine substitution converts the distal D4 cluster to an unprecedented modified non-cubane [3Fe-4S] cluster with a terminal sulfur ligand. Phylogenetic analysis revealed that the distal D4 non-cubane cluster is disrupted in most BamD homologues **(Extended Data Fig. 9d; Extended Data Table 2)**.

### Cryo-ET identifies electron-transferring flavoprotein as high-potential electron acceptor

To identify the high-potential electron acceptor and to clarify the cellular localisation and electron transfer networks, we examined the BCRII complex *in situ* with cryo-ET. We vitrified *G. metallireducens* cells grown with benzoate and nitrate, thinned cells by focused ion beam milling **(Extended Data Fig. 10a-d)**, collected tilt-series, and reconstructed tomograms of the native cellular interior. Analysis of these tomograms using template matching revealed that BCRII is dispersed throughout the cytoplasm rather than associated with the cytoplasmic membrane, as was reported before by immuno-electron microscopy **(Fig. 4a-c)**. This indicates that electron transfer to the MK pool does not occur via direct membrane association of BCRII, as previously assumed, and must instead involve a soluble electron carrier.

**Fig. 3.**
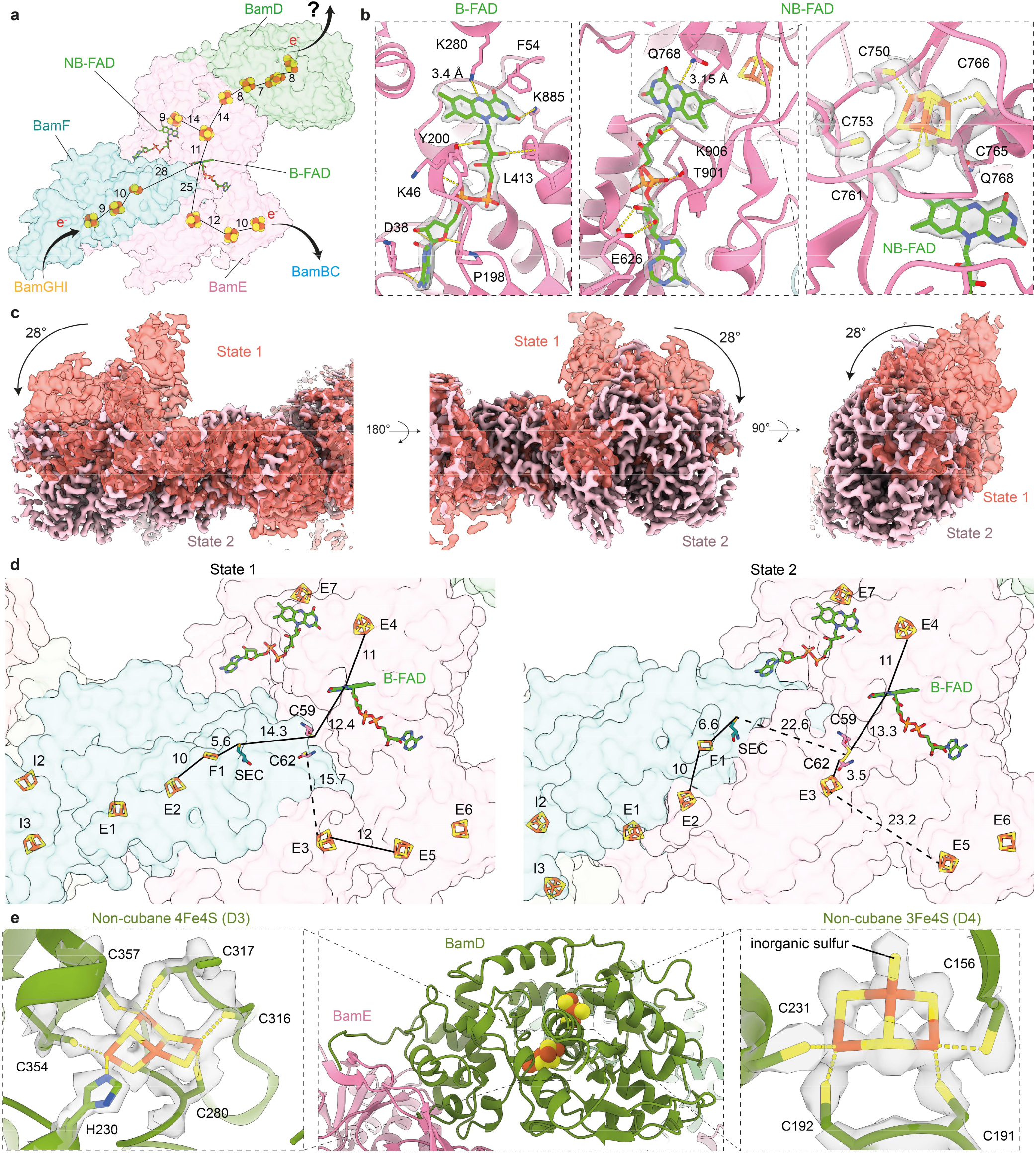
The BamE subunit mediates an HdrA-type electron bifurcation. **a**, Transparent surface representation of the BamDEF subcomplex showing all Fe–S clusters, FAD and selenocysteine (SEC). BamD contains one canonical non-cubane [4Fe–4S] (D3), and one non-cubane [3Fe– 4S] (D4) cluster. Electrons from BamGHI via confurcation reach BamE, where bifurcation directs them to the BamB and BamD branch. **b**, Enlarged view of BamE FADs with cryo-EM density. B-FAD is coordinated by Lys280 (∼3.4 Å to N5 of isoalloxazine, forming H-bond); NB-FAD by Gln768 (∼3.15 Å). **c**, Cryo-EM densities of two BCR II conformational states (1 and 2) showing ∼28° rotation of the peripheral arm; BamGHIF and BamE segments interfacing with BamF/C undergo major rearrangements. **d**, Cofactor distances in BamEF: in state 1, E3 is ∼16 Å and the selenocysteine ∼14 Å from Cys59–Cys62; in state 2, E3 approaches to ∼3.5 Å from the cysteine pair (forming disulfide), while selenocysteine moves to ∼23 Å. **e**, Cartoon of BamD showing the D3 [4Fe–4S] cluster coordinated by five cysteines and one histidine, and the D4 [3Fe–4S] cluster with terminal sulfur ligand coordinated by four cysteines. All distances in Å.

**Fig. 4.**
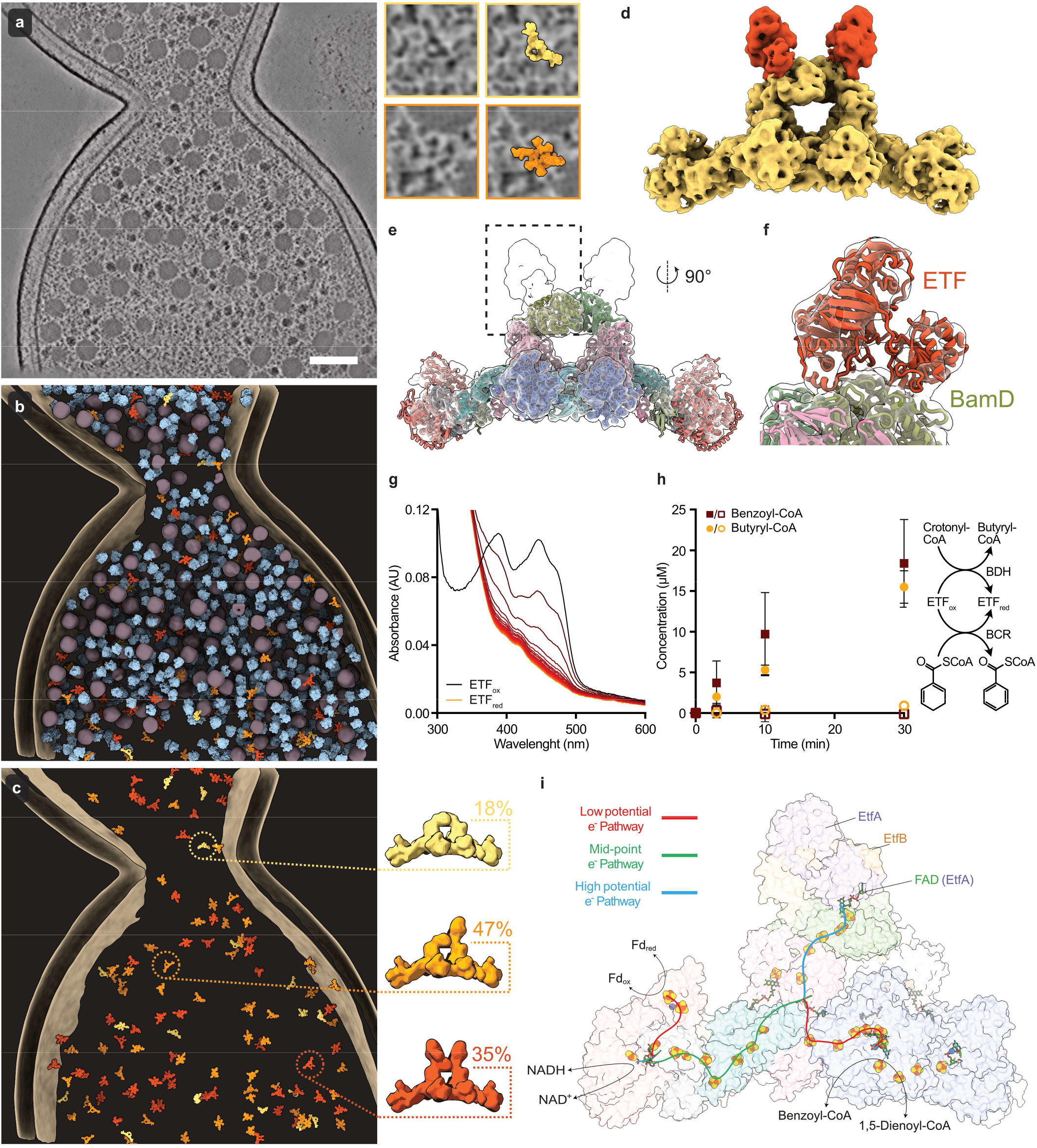
Identification of an ETF as the high-potential acceptor of BCRII. **a**, Slice through a cryo-electron tomogram of a *Geobacter metallireducens* cell, with side panels showing two BCRII complexes and mapped-back particles (slice thickness 7.64 Å; scale bar, 100 nm). **b**, Segmentation of the tomogram: BCRII in yellow/orange/red, ribosomes in light blue, storage granules in dark grey, membranes in tan. **c**, BCRII distribution and ETF occupancy: yellow is 0× ETF; orange is 1×; red is 2×; inset shows overall abundances from all analysed tomograms. For more examples, see **Extended Data Fig. 12. d**, Subtomogram average of BCRII, with Bam[(BC)_2_DEFGHI]_2_ in yellow and additional *in situ* density in red. **e**, Fit of single-particle Bam[(BC)_2_DEFGHI]_2_ model into the subtomogram average, highlighting additional density adjacent to BamD. **f**, Zoomed local map showing density consistent with ETF binding; model fitted with AlphaFold3 and Isolde^39^. **g**, Representative UV/vis spectra showing ETF (10 µM) reduction by BCR (0.1 µM) using 1,5-dienoyl-CoA (150 µM); control without BCRII in **Extended Data Fig. 13a. h**, Discontinuous assay coupling ETF (0.2 µM) reduction by BCRII (50 nM) with crotonyl-CoA (250 µM) conversion to butyryl-CoA via BCD (0.2 µM); filled/unfilled symbols indicate presence/absence of BCRII. Samples quenched at 0, 3, 10, 30 min and analysed by UPLC; unprocessed data in **Extended Data Fig. 13b**,**c. i**, Proposed electron transfer model: electrons from reduced ferredoxin and NADH enter via BamGHI, confurcate, then bifurcate at BamE FAD. High-potential electrons flow through BamD to ETF, which after reduction shuttles electrons to MK, while low-potential electrons reduce benzoyl-CoA at the Bam(BC)_2_ catalytic module.

To resolve the native structure of BCRII *in situ*, we applied an iterative subtomogram averaging workflow **(Extended Data Fig. 10e)**, yielding a 7.9 Å reconstruction of the complex. In parallel, ribosomes were resolved to 4.1 Å and their dimeric (100S) state to 8.8 Å **(Fig. 4d; Extended Data Fig. 11)**, enabling improved tilt-series refinement and providing structural context within the cellular environment.

Overall, the subtomogram average closely resembles the single-particle cryo-EM structure. Strikingly, we observed additional protein density adjacent to the two non-cubane Fe–S clusters of BamD, precisely at the terminus of the high-potential electron transfer pathway **(Fig. 4d**,**e)**. Focused 3D classification of this region yielded a local map with improved features, revealing a density consistent with an electron transferring flavoprotein (ETF) bound to BamD **(Fig. 4f; Extended Data Fig. 11e)**. Fitting ETF into the subtomogram average reveals that its domain II lies directly adjacent to the two non-cubane Fe–S clusters of BamD, positioning the α-FAD to receive the high-potential electron generated at the bifurcating FAD in BamE **(Fig. 4f**,**i)**. *G. metallireducens* encodes nine ETFs. One of these, BamOP, is upregulated during growth on benzoate^36^ and has a redox potential of around E°’ = +16 mV^37^. Subtomogram classification revealed that ETF is bound to approximately 59 % of BamD sites **(Fig. 4c; Extended Data Fig. 12)**, suggesting a transient interaction. After reduction, it likely dissociates and then regenerates by reducing MK via benzoate-induced ETF:MK oxidoreductase (EMO), a membrane-bound oxidoreductase that usually links fatty acid β-oxidation to the MK-pool in most MK-containing bacterial and archaeal lineages^38^.

Our results suggest the hierarchical coupling of a flavin-based electron confurcation, with Fd_red_ and NADH as low- and high-potential electron donors, to an FBEB event, with benzoyl-CoA and ETF as low- and high-potential electron acceptors. Attempts to measure such a coupled confurcation/bifurcation *in vitro* were hampered by short-circuit reactions from Fd_red_ to ETF (1.77 ± 0.57 µM min^-1^) **(Extended Data Fig. 13a)**. Even the use of Fd_red_ (via KGOR, α-ketoglutarate and CoA) and ETF (via acyl-CoA dehydrogenase and butyryl-CoA) regenerating systems could not circumvent these short-circuit reactions **(Extended Data Fig. 13b**,**c)**. To demonstrate the connection between high- and low-potential electron output modules, we used 1,5-dienoyl-CoA as an electron donor and assayed reduction of ETF alone **(Fig. 4g; Extended Data Fig. 13d)** or ETF in combination with butyryl-CoA dehydrogenase to reduce crotonyl-CoA **(Fig. 4h; Extended Data Fig. 13e-l)**. In both cases, BCRII-dependent electron transfer from 1,5-dienoyl-CoA to ETF was observed (k_cat_ = 132.6 ± 46.5 min^-1^ per BamD with excess ETF), providing direct *in vitro* evidence that ETF indeed serves as the second, high-potential electron-output module.

## Conclusion

The canonical strategy to overcome the conventional lower-potential redox limit of biological electron transfer, defined by the Fd_ox/red_ couple, relies on coupling to stoichiometric ATP hydrolysis. However, this mechanism is incompatible with the energetic constraints of strictly anaerobic microorganisms, such as sulfate-, Fe(III)-, or CO_2_-respiring bacteria that dominate marine and freshwater sediments and play central roles in global elemental cycles and bioremediation of aromatic pollutants.

A key conceptual advance is that BCRII assembles pre-existing electron-confurcating and -bifurcating modules in series and couples them to distinct electron input (Fd_red_, NADH) and output (benzoyl-CoA, ETF) units thereby enabling redox chemistry otherwise inaccessible to these organisms. This modular recombination of unrelated redox components exemplifies how evolution expands metabolic capacity without introducing fundamentally new mechanisms. In BCRII, the low- and high-potential output branches span an exceptional ∼640 mV gap, enabling the thermodynamically demanding reduction of aromatic rings beyond the conventional biological redox window as follows:

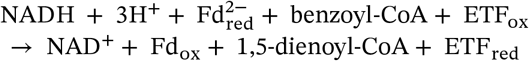

Among characterised FBEB systems, BCRII is exceptional in using Fd_red_, together with NADH, as electron donor, whereas in all other reported FBEB enzymes, Fd functions as the low-potential acceptor. The identification of ETF as the physiological high-potential acceptor positions BCRII catalysis within the cellular redox network, linking the degradation of aromatic compounds to downstream electron transfer. Within this framework, low-potential electrons are directed towards aromatic ring reduction, while high-potential electrons feed into the respiratory chain via ETF and ETF:MK oxidoreductase.

The BCRII system illustrates how redox networks can be organised into large, multi-enzyme assemblies that integrate electron transfer, storage, and gating within a single functional unit. BCRII thus represents a prototypical redox assembly in which discrete input and output modules are functionally coupled with energy-coupling confurcation and bifurcation units. This framework highlights nature’s combinatorial potential in cellular redox chemistry and offers a blueprint for engineering synthetic, modular electron-transfer networks.

## Materials and Methods

### Bacterial strains and plasmids

*G. metallireducens* GS-15 (DSMZ 7210) and *Thauera aromatica* K172 (DSMZ 6984) were used for wild type purifications. *E. coli* BL21 (DE3) was used to heterologously produce the ETF (Gmet_2153/2152) from *G. metallireducens* with a C-terminal twin-strep tag in a pPR-IBA101 vector as previously described^37^. *E. coli* BL21 (DE3) was also used to produce Glutaryl-CoA dehydrogenase (GDH, Gmet_2075) from *G. metallireducens* with a C-terminal 6xHis-tag in a pEXP5-CT/TOPO vector^40^. *E. coli* MC4100 was used to heterologously produce BamGHI, (Gmet_2079-2081), after cloning of the genes into a pCE vector with an N-terminal Strep-Tag. *E. coli* C43 (DE3) was used to heterologously produce an acyl-CoA dehydrogenase (BDH, desgi_1918) from *Desulfotomaculum gibsoniae*, which was cloned into a pET28b vector with a C-terminal 6×His-tag.

### Cultivation of cells

*G. metallireducens* was cultivated anaerobically in a 200 L fermentor at 30 °C in a mineral salt medium as previously described^41^ with benzoate (1-5 mM) serving as the sole carbon source and nitrate (3 mM) as electron acceptor. Cells were harvested by centrifugation (20,000×*g* at 4 °C) and stored in liquid nitrogen. *G. metallireducens* cells used for cryo-ET were cultivated anaerobically in the same medium in 100 mL bottles.

*E. coli* BL21 (DE3)/pPR_IBA101_ETF, *E. coli* BL21 (DE3)/pEXP5-CT/TOPO_GDH and *E. coli* C43 (DE3)/pET28b_BDH were cultivated in 2 L shaking flasks in auto-inducing medium (1 % [w/v] yeast extract, 1 % [w/v] tryptone, completed with 0.2 % [w/v] D-lactose, 0.05 % [w/v] glucose, 0.5 % (w/v) glycerol, 25 mM Na_2_HPO_4_, 25 mM KH_2_PO_4_, 50 mM NH_4_Cl, 5 mM Na_2_SO_4_ and 2 mM MgSO_4_) with 100 ng mL^−1^ ampicillin (ETF and GDH) or 75 ng mL^−1^ kanamycin (BDH). The cultivation was started at 37 °C and 160-180 rpm for 4 h and followed by an overnight incubation at 20 °C at 140 rpm. Cells were harvested by centrifugation (6,000 × *g* at 4 °C) and stored at −20 °C.

*E. coli* MC4100/pCE_BamGHI was cultivated under anaerobic conditions in buffered terrific broth medium at pH 7.5 (50 mM fumaric acid, 12 g L^−1^ tryptone, 24 g L^−1^ yeast extract, 2.5 g L^−1^ glycerol, 0.76 mM Fe-(III)-citrate, 15 mM KH_2_PO_4_, 74 mM K_2_HPO_4_, 6.5 g L^−1^ KOH). Cultivation was performed in either 2 L bottles with rubber stoppers or a 200 L fermentor. Following inoculation, the headspace was flushed with N_2_ gas and the cells were grown at 37 °C. BamGHI production was induced in the exponential growth phase with 0.2 % arabinose and the cells were supplemented with 3 mM sodium nitrate, 0.8 mM magnesium sulphate, 0.1 mM calcium chloride, trace elements and vitamin solution. Additionally, the incubation temperature was lowered to 30 °C for approximately 18 h and then to 20 °C for 4 h. Cells were harvested by centrifugation (6,000 × *g* at 4 °C for bottle cultivation, 20,000 × *g* at 4 °C for fermentor cultivation) and stored in anaerobic rubber-stoppered bottles at −80 °C.

### Enrichment of proteins

### Bam[(BC)_2_DEFGHI]_2_ complex

The enrichment of the BCRII complex from *G. metallireducens* was adapted from the method described by Huwiler^10^. All steps were carried out under strictly anaerobic conditions under a N_2_/H_2_ atmosphere (95 %/5 %). For the preparation of crude extracts, 10-30 g of cells were suspended in buffer A (50 mM potassium phosphate, 5 mM MgCl_2_, 5 µM FMN; pH 6.5; 1.5 mL buffer g^−1^ cells), containing 100 mM KCl, 10 mM dithioerythritol (DTE) and a spatula tip of DNase I and RNase A, respectively. *G. metallireducens* cells were lysed by stirring for 1 h at 4 °C with 1.5 mg lysozyme per g cell mass. The crude extracts were centrifuged at 200,000 × *g* (1 h, 4 °C) and the pellet was washed in buffer A with 100 mM KCl and centrifuged again, after which the supernatants were combined. The supernatants were filtered and applied to a DEAE Sepharose FF column (Cytiva; 40 mL, 2.6 cm diameter, 2 mL min^−1^ flow rate), equilibrated with buffer A. Unspecifically bound proteins were washed from the column with two to three column volumes of the respective buffer containing 200 mM KCl. The DCO activity containing BCRII eluted at 350 mM. The protein eluted in a volume of approx. 50 mL and was concentrated to 0.5-1 mL using 100,000 MWCO centrifugal filters (Vivaspin® Turbo, Sartorius). The concentrated fractions were applied to a Superose™ 6 Increase 10/300 GL column (Cytiva; flow rate 0.5 mL min^−1^), equilibrated with buffer A without FMN, containing 100 mM KCl. 1,5-dienoyl-CoA:methyl viologen oxidoreductase (DCO) activity assigned to BCRII eluted in a brownish high molecular weight fraction between 12-14 mL. The fractions were concentrated to at least 10 mg mL^−1^ and typically used the next day without freezing, but stored in anaerobic glass vials overnight at 4 °C. For structure determination by cryo electron microscopy (cryo-EM), the BCRII preparations were concentrated to at least 20 mg mL^−1^ and stored/transported under strictly anoxic conditions at <8 °C. The enzyme was diluted to the final concentration immediately before grid preparation, also carried out in a N_2_/H_2_ atmosphere (95 %/ 5%).

### Enrichment of BamGHI

Heterologously produced BamGHI from *G. metallireducens* was enriched under a N_2_/H_2_ atmosphere (95 %/5 %) using Strep-Tag affinity chromatography. Cells were suspended in buffer C (50 mM HEPES/KOH, 150 mM NaCl, 5 µM FMN; pH 7.6) with lysozyme (1 mg g^−1^ cells) and spatula tips of DNase I and RNase A. Following cell lysis using a French pressure cell at 1,100 psi and centrifugation at 200,000 × *g* (1 h, 4 °C), the supernatant was loaded to a 5 mL Strep-Tactin®XT 4Flow® high capacity FPLC column (IBA GmbH; 3 mL min^−1^ flow rate), pre-incubated with buffer C. BamGHI was eluted using buffer C with 50 mM D(+)biotin and concentrated to at least 5 mg mL^−1^ using 50,000 MWCO centrifugal filters. The sample was frozen in liquid nitrogen and stored under anoxic conditions at –80 °C.

### Enrichment of ETF

Heterologously produced ETF from *G. metallireducens* was enriched using Strep-Tag affinity chromatography, as previously described^37^. Cells were suspended in buffer D (50 mM HEPES/KOH, 150 mM NaCl, pH 7.6) with a spatula tip of DNase I. A French pressure cell (1,100 psi) was used to lyse the cells and the lysate was centrifuged at 200,000 × *g* (1 h, 4 °C). The supernatant was applied to a 5 mL Strep-Tactin®XT 4Flow® high capacity FPLC column (3 mL min^−1^ flow rate), equilibrated with buffer D. ETF was eluted using buffer D with 50 mM D(+)biotin and concentrated to at least 50 mg mL^−1^ using 50,000 MWCO centrifugal filters. The enzyme was frozen in liquid nitrogen and stored at –80 °C in buffer D with 20 % (v/v) glycerol.

### Enrichment of butyryl-CoA dehydrogenase (BDH)

Heterologously produced BDH from *D. gibsoniae* was enriched by His-tag affinity chromatography. Cells were suspended in buffer E (20 mM tricine, 250 mM NaCl, 10 % glycerol; pH 7.4) with a spatula tip of DNase I. After cell lysis using a French pressure cell (1,100 psi) and centrifugation at 200,000 × *g* (1 h, 4 °C), the supernatant was applied to a 5 mL HisTrap™ chromatography column (Cytiva; 3 mL min^−1^ flow rate) pre-incubated with buffer E. Weakly interacting proteins were washed from the column using buffer E containing 40 mM imidazole. BDH was eluted with buffer E containing 400 mM imidazole and concentrated to at least 20 mg mL^−1^ using 30,000 MWCO centrifugal filters. The enzyme was desalted (PD-10 desalting column; GE Healthcare), frozen in liquid nitrogen and stored at –80 °C.

### Enrichment of glutaryl-CoA dehydrogenase (GDH)

Heterologously produced GDH from *G. metallireducens* was enriched by His-tag affinity chromatography. Unless specified, steps were carried out as described for BDH. During GDH enrichment, buffer F (20 mM Tris/HCl, 250 mM KCl, 10 mM imidazole, 5 % glycerol; pH 7.8) was used for cell suspension and to equilibrate the column. GDH was eluted with a linear gradient of buffer F containing 0-500 mM imidazole as described^40^.

### Enrichment of Fd

Fd from *G. metallireducens* was enriched in a N_2_/H_2_ atmosphere (95 %/5 %) as previously described^10^. For the preparation of crude extract, 10 g cells were suspended, lysed and centrifuged analogous to BCRII enrichment. The 200,000 × *g* supernatant was applied to a DEAE Sepharose FF column (40 mL, 2.6 cm diameter, 2 mL min^−1^ flow rate), equilibrated with buffer A at pH 7.2. The column was washed with buffer A (pH 7.2) containing 200 mM KCl. The fraction eluting over 25 mL in a subsequent step with 300 mM KCl had a brownish colour. The fraction was concentrated to 0.5 mL using 10,000 MWCO centrifugal filters and applied to a Superdex75 Increase column (24 mL, 10/300 GL, 0.5 mL min^−1^ flow rate) equilibrated with buffer A (pH 7.2) containing 100 mM KCl. The low molecular weight fraction with a dark brown colour eluted over 1.5 mL. A molar extinction coefficient of ε_390nm_ = 30,000 M^−1^ cm^−1^ was used to determine the concentration of Fd. The A_390 nm_/A_280 nm_ ratio indicating protein purity was 0.76, consistent with similar proteins, where a ratio of 0.75 was determined^42^.

### Enrichment of α-ketoglutarate:Fd oxidoreductase (KGOR)

The KGOR from *T. aromatica* cells grown on benzoate and nitrate was enriched in a N_2_/H_2_ atmosphere (95 %/5 %) as described^43^. For preparation of cell extracts, 30-50 g cells were suspended in buffer G (20 mM triethanolamine-HCl (pH 7.8), 4 mM MgCl_2_, 10 % glycerol, 0.2 mM thiamine diphosphate [TPP], and 1 mM DTE). After cell lysis using a French pressure cell (1,100 psi) and centrifugation at 200,000 × *g* (1 h, 4 °C), the supernatant was applied to a DEAE Sepharose FF column (40 mL, 2.6 cm diameter, 2 mL min^−1^ flow rate) equilibrated with buffer G. KGOR activity eluted with buffer G containing 90 mM KCl in a volume of 100 mL. The eluate was applied to a Q-Sepharose column (50 mL, 2.6 cm diameter, 2 mL min^−1^ flow rate) equilibrated with buffer G containing 90 mM KCl and KGOR activity eluted in the flow through (approx. 120 mL). The KGOR activity-containing fractions were applied to a Reactive-Red-agarose (120 Type 3000-CL, Sigma Aldrich; 25 mL 2.6 cm diameter, 3 mL min^−1^ flow rate) column equilibrated with buffer G. The column was washed with two bed volumes of buffer G, two bed volumes of buffer G plus 2 mM α-ketoglutarate and 500 mM KCl, and two bed volumes of buffer G plus 2 mM α-ketoglutarate and 1.2 M KCl. KGOR eluted in the last step and was concentrated to 1 mL using 50,000 MWCO centrifugal filters. The concentrated protein was applied to a Superdex 200 gel filtration column (300 mL; diameter, 2.6 mL, 2 mL min^−1^ flow rate) in buffer G with 100 mM KCl. KGOR was concentrated to at least 4 mg mL^−1^, frozen in liquid nitrogen and stored anaerobically at –80 °C.

### Enzyme assays

All enzyme assays were performed in a glove box under anoxic conditions in a N_2_/H_2_ atmosphere (95 %/5 %) at 30 °C. A typical assay was performed in a 50 mM potassium phosphate buffer (pH 7.0) with 100 mM KCl, 5 mM MgCl_2_ and 2 mg mL^−1^ BSA. UV/visible spectroscopic data were acquired using a UV-1800 spectrophotometer (Shimadzu) with quartz cuvettes (1 cm diameter). Analysis was performed using UVProbe software, version 2.43 (Shimadzu), and spectra were depicted using Prism 6 (GraphPad).

### DCO and NOR assays

The 1,5-dienoyl-CoA:methyl viologen (MV, DCO)^10^ and NADH:benzyl viologen (BV) oxidoreductase (NOR)^23^ activities were determined spectroscopically as described in 200-400 µL. The DCO activity assays contained 0.1 mM 1,5-dienoyl-CoA and 0.5 mM MV and the time dependent MV reduction was followed at 730 nm (Δε = 2,400 M^−1^ cm^−1^). The NOR activity was determined with 0.5 mM NADH and 1 mM BV at 600 nm (Δε = 7,400 M^−1^ cm^−1^). Both types of activity assays were started with 20-60 µM BCRII.

### Reduced methyl viologen (MV_red_):NAD^+^ oxidoreductase assay

The MV_red_-dependent NAD^+^ reduction activity was determined spectroscopically. The assay mixture (200-400 µL) contained 1 mM MV_red_ (previously reduced with sodium dithionite), 0.5 mM NAD^+^ and was started with 20-50 µM BCRII. The time-dependent oxidation of MV was monitored at 730 nm (Δε = 2,400 M^−1^ cm^−1^).

### ETF reduction with 1,5-dienoyl-CoA via BCRII

The kinetics following the reduction of ETF were measured at 445 nm (Δε = 14,000 M^−1^ cm^−1^)^37^ with 20 µM ETF, 25-50 nM BCRII, and 0.3 mM 1,5- dienoyl-CoA. Reduction of ETF was also monitored spectroscopically from 800 to 300 nm (all spectra were normalised for their absorbance at 800 nm) with 10 µM ETF, 50-100 nM BCRII and 0.15 mM 1,5-dienoyl-CoA. Spectra were taken every 60 s. Controls did not contain BCRII.

Discontinuous assays were performed and followed by ultra-performance liquid chromatography (UPLC) analysis. In this setup, the reduction of ETF by 1,5-dienoyl-CoA via BCRII was coupled to the reduction of crotonyl-CoA via BDH. The assay mixture contained 0.25 mM 1,5-dienoyl-CoA, 0.25 mM crotonyl-CoA, 50 nM BCRII, 0.2 µM ETF and 0.2 µM BDH. Controls were performed without BCR. Samples of 20 µL were taken at 0, 3, 10 and 30 min and stopped by addition of 40 µL of methanol. After centrifugation, the supernatants were analysed on an Acquity UPLC® device (H-Class, Waters GmbH) using an Acquity UPLC HSS T3 column (2.1 mm × 100 mm, 1.8 µm particle size) with an acetonitrile/potassium phosphate (pH 7) gradient of 0-20 % acetonitrile over 5 min. The CoA esters were distinguished by their retention times and characteristic spectra, which were compared to standards. Standards were prepared and measured analogously to samples. Formation of benzoyl-CoA was corrected for the formation of cyclohex-1-ene-1-carbonyl-CoA (Ch1CoA), since the BCRII catalyses the disproportionation of 1,5-dienoyl-CoA to these two products^16^.

### Reduction of BamGHI with reduced ferredoxin (Fd_red_) and NADH

The reduction of BamGHI was followed by absorption spectroscopy (800-300 nm), and all spectra were normalised for their absorbance at 800 nm. BamGHI (0.36 µM) was reduced for 30 min either by 0.1 mM NADH or by 0.25 mM CoA, 0.33 mM α-ketoglutarate, 0.4 µM Fd and 0.3 µM KGOR. Spectra were taken every 60 s and corrected for dilution, as well as for the absorbance of Fd and KGOR. For better comparison, all spectra were normalised to the same absorbance at 310 nm prior to the addition of the electron donors. Reduction with 6 µM sodium dithionite was set as 100 % reduction. The extent of reduction per donor was determined at 450 nm. BamGHI reduction by NADH was normalised to the FMN occupancy of BamH.

### Electron bifurcation and confurcation assays

The combined electron confurcation/bifurcation assays with BCRII were conducted in various conditions. The electron donors used were NADH and Fd_red_. NADH was used in concentrations of 0.3-2 mM and regenerated using 0.2-2 U mL^−1^ mM formate DH, 1-5 mM formate and 0.2-1 mM NADH. The concentrations of the Fd_red_ electron donor system were 1-8 µM Fd, 0.2 µM KGOR, 0.5-2 mM CoA and 0.5-2 mM α-ketoglutarate. The electron acceptors were 50-400 µM benzoyl-CoA and ETF. ETF was either used at 15-300 µM or at 15-40 µM ETF regenerated by 2-10 µM BDH and 0.2-1 mM crotonyl-CoA. Assays were conducted either with 0.4-2 µM BCRII, or with desalted cell extract from *G. metallireducens*. Assays with enriched BCRII were also performed in the presence of 50 µM FMN and FAD or with pre-reduced and subsequently desalted BCRII using either sodium dithionite or 1,5-dienoyl-CoA as electron donors. Furthermore, the electron confurcation/bifurcation assays were also performed with *G. metallireducens* raw extracts thereby attempting to regenerate potentially reduced MK by nitrate reductase using 2-3 µM sodium nitrate as terminal electron acceptor in the presence or absence of 1 mM Mg ATP or 5 mM DTT.

The reverse BCRII reaction was tested with 50-200 µM 1,5-dienoyl-CoA and reduced ETF as electron donors. The ETF reduction system was 0.4-15 µM ETF, 0.2-2 µM GDH and 0.05-2 mM glutaryl-CoA. The concentrations of the electron acceptors were 0.1-0.15 µM NAD^+^ and 30 µM Fd. The assays were conducted with 0.05-1.25 µM BCRII from *G. metallireducens*.

All assays were also conducted under different pre-incubation conditions of BCRII or cell extracts with the electron donors or acceptors.

To determine the 1,5-dienoyl-CoA formation from benzoyl-CoA or 1,5-dienoyl-CoA-dependent NADH formation, assays were analysed using an Acquity UPLC® device (H-Class, Waters GmbH) using an Acquity UPLC HSS T3 column (2.1 mm × 100 mm, 1.8 µm particle size). An acetonitrile/potassium phosphate (pH 7) gradient of 0-20 % acetonitrile over 5 min was used to separate the assay components. Alternatively, a gradient of 2-10 % acetonitrile over 1.5 min followed by 10-30 % acetonitrile over 4.5 min was employed. The compounds were distinguished by their retention times and spectra, which were compared to standards.

### KGOR, BDH and GDH activities

The activities of KGOR, BDH and GDH were tested spectroscopically. KGOR activity to reduce Fd was validated in assays containing 1.7-2.3 µM KGOR, 6-14 µM Fd, 0.4-1 mM CoA, which were started by the addition of 0.5-1 mM α-ketoglutarate. The activity of BDH to regenerate oxidised ETF was controlled with 1 µM BDH, 10-20 µM ETF (pre-reduced with sodium dithionite and desalted) and started with 0.1 mM crotonyl-CoA. GDH activity to reduce ETF was tested with 0.2-1 µM GDH, 10 µM ETF and started with 0.2-0.5 mM glutaryl-CoA.

### Short circuit reactions during electron bifurcation assays

Short circuits between electron donors and ETF in electron bifurcation assays were measured at typical concentrations. The reduction of ETF was spectroscopically monitored at 445 nm (Δε = 14,000 M^−1^ cm^−1^)^37^ using a concentration of 20 µM ETF. Furthermore, the assay mixture either contained 0.5 mM NADH, or 0.33 mM CoA, 0.33 mM α-ketoglutarate, 0.3 µM KGOR and 1 µM Fd. The reaction was started by the addition of NADH or Fd, respectively.

### Determination of metal cofactors and flavin occupancy

Iron contents were determined colorimetrically, as described^44^. FMN and FAD content was determined by precipitating a 30 µL protein sample (1-5 mg mL^−1^) with 5 µL of 1 M H_2_SO_4_ and incubating for 15 min. After centrifugation at 13,000 × *g*, the supernatant was collected and the pellet was washed with 20 µL of 20 mM H_2_SO_4_ and centrifuged again. Both supernatants were combined and analysed by an Acquity H-Class UPLC® device (Waters GmbH) with an Acquity UPLC HSS T3 column (2.1 mm × 100 mm, 1.8 µm particle size). The spectra and retention times were compared with those of the standards. All protein concentrations were routinely determined by the Bradford method, using BSA as the standard.

### Protein identification by mass spectrometry

Protein purity was analysed by sodium dodecyl sulfate polyacrylamide (SDS) gel electrophoresis and protein bands were identified by mass spectrometry (MS) of tryptic peptides, as previously described^9^. An Acquity I-class UPLC® device (Waters) and a Waters Peptide CSH C18 column (2.1 mm × 150 mm, 1.7 µm particle size) were used to separate the peptide fragments with a gradient of 1-40% ACN/0.1 % formic acid (v/v) in water/0.1 % formic acid (v/v) (flow rate 0.1 mL min^−1^). MS analysis was performed on a Waters Synapt G2-Si HDMS ESI/Q-TOF system in positive HD-MS^E^ mode (3 kV cone voltage, 120 °C source temperature, 400 °C desolvation temperature, 800 L h^−1^ N_2_ desolvation gas flow, 6.0 bar nebuliser pressure). Fragment detection and analysis using the ProteinLynx Global Server (Waters) were performed by matching with the UniProt database of *G. metallireducens* (minimal fragment ion matches per peptide = 3, minimal fragment ion matches per protein = 7, minimal peptide matches per protein = 1, maximal false discovery rate 4 %) as described^45^

### Cryo-EM grid preparation

For cryo-EM data acquisition, Quantifoil R2/1 copper grids (300 mesh; Quantifoil Micro Tools, Germany) were glow-discharged for 25 s at 15 mA using a PELCO easiGlow device (Ted Pella, USA). Prior to vitrification, 30 mg mL^−1^ of highly concentrated protein was incubated with 500 µM NADH and pre-reduced Fd (5 µM Fd, 0.2 µM KGOR, 1 mM α-ketoglutarate, 1 mM CoA) to establish physiologically relevant reducing conditions and capture potential redox-driven conformational states. The sample was subsequently diluted to 1.5 mg mL^−1^, and 4 μL was immediately applied to the grids and vitrified. Vitrification was performed using a Vitrobot Mark IV (Thermo Fisher Scientific, USA) with a blot force of 7 for 7 s at 4 °C and 100 % humidity, followed by plunge-freezing in liquid ethane. All sample preparation and handling steps were carried out under strictly anaerobic, redox-controlled conditions in a Coy vinyl anaerobic chamber (Coy Laboratory Products, USA) containing 95 % N_2_ and 5 % H_2_.

### Cryo-EM data collection, processing, and model building

Cryo-EM data were collected using Smart EPU software on a Titan Krios G4 transmission electron microscope (Thermo Fisher Scientific, USA) operating at 300 keV and equipped with a Falcon K4i direct electron detector. Images were recorded at a nominal magnification of 165,000 ×, corresponding to a calibrated pixel size of 0.732 Å. Data were acquired in counting mode with a total exposure of 60 e− Å^−2^ and stored in EER format. A total of 21,619 micrographs were collected. All data processing was performed in cryoSPARC^46^. EER movies were fractionated into 60 frames, followed by patch motion correction^47^ and CTF estimation. An initial set of particles was manually picked from 200 micrographs and used to iteratively train a Topaz model^48^ on a subset of 500 micrographs until the optimal model was generated. The trained model was then applied to the full dataset, yielding approximately 864,000 particles, which were extracted with a box size of 480 pixels 4 × binned and subjected to *ab initio* reconstruction. One major class (536,089 particles) displaying the characteristic Ω-shaped core with diffuse arms was selected for heterogeneous refinement. Subsequent classification yielded four classes, of which one (263,000 particles) exhibited well-defined features of the BCRII complex. This class was further refined using hard classification, resulting in three subclasses: one containing only the core, and two containing both the core and peripheral arms. One subclass displayed both arms (with asymmetric density), whereas the other showed only the left arm. Particles from the two arm-containing classes were combined, re-extracted un-binned with a box size of 588 pixels, and subjected to non-uniform (NU) refinement, followed by reference-based motion correction and a second round of NU refinement. Refinement with C1 and C2 symmetry yielded maps at 2.52 Å and 2.3 Å resolution (FSC = 0.143), respectively. The C2 map prominently resolved the core (BamEDBC) region, whereas the C1 map showcased the core including the arm (BamGHIF) of the left side of the complex. To further improve the core reconstruction, a soft mask was applied to the core (BamEDBC), followed by masked 3D classification without alignment. Particles from four of five classes (150,032 particles) were selected and refined to 2.0 Å resolution. Symmetry expansion (C2; 302,064 particles) and local refinement of a single BamED protomer further improved the resolution to 1.9 Å. Additional local refinement with a focused mask on the BamD dimer yielded a 2.0 Å map. For the BamBC region, C4 symmetry expansion (604,128 particles) followed by masked local refinement produced a map at 1.97 Å resolution. The left arm (Arm 1) was resolved using the C1-refined map. A mask was applied to the BamGHIF region, followed by 3D classification into five classes, revealing two distinct conformational states. State 1 (two classes; 40,472 particles) was refined to 2.55 Å, whereas State 2 (one class; 15,799 particles) was refined to 3.0 Å. Focused refinement of the BamED region within these states yielded resolutions of 2.3 Å (State 1) and 2.5 Å (State 2). The right arm (Arm 2) was reconstructed from a separate subclass obtained during heterogeneous refinement. Masked 3D classification without alignment of the BamGHIF region produced five classes, of which two were refined to 3.5 Å resolution. A weighted composite map was generated by combining locally refined maps using PHENIX combine focused maps^49^.

Initial models for all subunits were generated using AlphaFold^50^ based on sequences from *Geobacter metallireducens* GS-15 (GenBank ABB32381.1, ABB32382.1). Models were fitted into cryo-EM densities using UCSF Chimera (v1.17.3) and manually adjusted in Coot (v0.9.8.92). Real-space refinement was performed iteratively in PHENIX (v1.21-5207). Cofactors and ligands were manually placed in Coot prior to refinement. Figures were prepared using UCSF ChimeraX (v1.6.1).

### Cell vitrification

*Geobacter metallireducens* GS-15 cultures were grown anaerobically in mineral salt medium with 1 mM benzoate and 3 mM nitrate at 30°C to an OD_600_ of 0.2. Cultures were centrifuged for 15 min at 1,900 × *g* to concentrate the cells to an OD_600_ of approximately 13. A 4.5 μL aliquot of the concentrated cell suspension was applied to holey carbon R2/1 or R1.2/1.3 200-mesh copper TEM grids (Quantifoil) under anaerobic conditions, and plunge-frozen in liquid ethane using a Vitrobot Mark IV (Thermo Fisher Scientific) (settings: blot force = –5, blot time = 7-10 s, temperature = 4° C). A Teflon ring was placed on the sample side to prevent cells from adhering to the filter paper during blotting. Samples were stored in liquid nitrogen until use.

### Cryo-focused ion beam (FIB) milling

Sample grids were clipped into Autogrids (Thermo Fisher Scientific) and subjected to cryo-focused ion beam milling in an Aquilos 2 dual-beam instrument (Thermo Fisher Scientific) using gallium ions at 30 kV accelerating voltage. Grids were loaded using the standard 35° pre-tilted holder and coated with an organometallic platinum layer using the gas injection system (20 s deposition).

Because cells formed an approximately continuous monolayer across the grid surface, virtually any position within a grid square could be targeted for lamella preparation. To exploit this feature, the milling angle was initially set to 90° relative to the plane of the grid (corresponding to a 45° holder pre-tilt), and each grid square was milled using a nominal beam current of 500 pA to generate 14-16 trench-milled positions.

After trench milling across the selected grid squares, the tilt angle was adjusted to achieve a milling angle of 11-18°. Lamellae within each grid square were then thinned in parallel using multiple cleaning cross-section patterns (for example, 16 lamellae required 32 patterns) with a nominal beam current of 100 pA. Lamella thickness was gradually reduced by decreasing the separation between the upper and lower milling patterns from 1200 nm to a final target thickness of approximately 180 nm (steps: 1200, 800, 400, 250 and 180 nm). See (Extended Data Fig. 10a-d) for representative lamellae and acquisition layout.

To minimise grid charging and stage drift, milling was performed continuously while the pattern distances were updated every 5-8 min without interrupting the process. This workflow enabled the production of approximately 100 lamellae within an 8-hour milling session.

### Cryo-electron tomography (cryo-ET) data acquisition

Cryo-ET datasets were acquired on a Titan Krios G4 transmission electron microscope (Thermo Fisher Scientific) equipped with a Falcon 4i direct electron detector and a Selectris X energy filter. The energy filter was operated with a 10 eV slit width and the zero-loss peak was tuned prior to each data acquisition session.

Tilt series were collected using a dose-symmetric tilt scheme implemented in Tomography 5 software (Thermo Fisher Scientific). The nominal tilt range was ±60°, with tilt increments of 2° or 2.5°. Acquisition was initiated at either +10° or −10° to compensate for the lamella milling angle. The target defocus was adjusted for each tilt series in steps of 0.3 μm over a range from −2 μm to −4.5 μm. Data were recorded in Falcon 4i EER mode at a nominal pixel size of 1.98 Å. The pixel size was later calibrated to 1.91 Å by fitting the BCRII subtomogram average to the atomic model of the BCRII complex. The electron dose per tilt ranged from 2.0 to 2.7 e^−^ Å^−2^ depending on the dataset.

### Cryo-ET data processing and subtomogram averaging

Tilt series data processing was primarily carried out using the WarpTools pipeline^51^ (see Extended Data Fig. 10e for an overview of the processing workflow). Raw EER frames were grouped to achieve an exposure of approximately 0.1-0.2 e^−^A^−2^ per group, followed by full-frame motion correction and CTF estimation on a 2 × 2 grid. Tilt series alignment was performed using AreTomo2^52^, and defocus handedness was evaluated using Warp-Tools and defocusgrad (https://github.com/CellArchLab/cryoet-scripts/tree/main/defocusgrad) and subsequently inverted. Tilt series CTF parameters were then estimated, and 3D-CTF corrected tomograms were reconstructed at 7.64 Å px^−1^.

Initial BCRII and ribosome picks were obtained by template matching with pytom-match-pick (Chaillet et al., 2023). The single-particle cryo-EM map of BCRII, masked around the Ω core of the complex, was used as the search template for BCRII particles. A bacterial 70S ribosome map (EMD-45266) served as the ribosome template. From a combined dataset of 63 tomograms, this yielded 5,675 BCRII and 10,186 ribosome picks. Subtomograms were reconstructed in WarpTools 3.82 Å px^−1^, and subjected to initial averaging in RELION 4.0.2^53^. To remove false positives, BCRII particles were subjected to iterative 3D classification without alignments (five classes, T = 4), resulting in a curated set of 3,698 particles.

The curated BCRII and ribosome particle sets were jointly imported into M^54^ for refinement of tilt series alignments, image-space warping and CTF estimation. The resulting reconstructions achieved global resolutions of 12 Å (BCRII) and 4.7 Å (ribosome). Using the refined parameters from M, optimised tomograms were reconstructed in WarpTools, and template matching was repeated using the initial BCRII and ribosome subtomogram averages as updated search templates. For BCRII, picks were thresholded using a cutoff determined automatically for each tomogram from the fitted maximum local correlation coefficient histogram. For tomograms with excessive false positives, extraction thresholds were determined using the tm_find_threshold script (https://github.com/CellArchLab/cryoet-scripts/blob/main/tm/tm_find_thresh), which estimates a cutoff from the sorted template-matching scores by detecting where the approximately linear high-score region transitions to the background distribution. This second round of template matching yielded higher correlation scores and increased the number of detected particles per tomogram. In total, 3,457 BCRII particles were picked from the 32 highest-quality tomograms. For ribosomes, a dual-thresholding strategy^55^, including tophat-filtered ribosome score maps, resulted in 17,026 ribosome particles from the same set of 32 tomograms.

As before, subtomograms were reconstructed in WarpTools at 3.82 Å px^−1^ and processed in RELION. BCRII false positives were removed by 3D classification (five classes, T = 2). The final curated sets of particles comprised 2,569 BCRII and 17,026 ribosomes. These were refined in RELION, resampled to 1.91 Å px^−1^, and imported into M. Refinement in M consisted of iterative cycles of particle pose, image warp (3x3) and defocus refinement. This was followed by an additional round of image warp (3x3), stage angles and particle pose refinement and one round of image warp (3x3), particle pose and CTF refinement. Per-series and per-tilt weights were then applied, and particle temporal trajectories refined. A final refinement cycle including image warp (6x6), stage angles, CTF, and particle poses yielded maps with global resolutions of 7.9 Å for BCRII and 4.1 Å for the ribosome (Extended Data Fig. 11a,d).

To assess ETF-binding site occupancy on BamD, the final BCRII particle set, as used for the final consensus refinement in M, was instead subjected to 3D refinement in RELION 4.0.2 with C2 symmetry imposed and subsequently symmetry-expanded. Focused 3D classification without alignments (two classes, T = 0.5), using a mask covering the ETF region, revealed ETF occupancy in approximately 59 % of sites. To reconstruct a focused map of the ETF-BamD interaction, symmetry-expanded particles corresponding to the ETF-bound state were re-centered on the interface and re-extracted at 1.91 Å px^−1^. Reconstruction in M without further refinement yielded a 10 Å map of the ETF-BamD region. Finally, to analyse the distribution of BCRII complexes with different ETF occupancies, a custom Python script was used to sort all BCRII particles into sets with 0, 1 or 2 occupied ETF-binding sites. The spatial distribution of these occupancy states was visualised by mapping the corresponding particle subsets back onto the tomograms in ArtiaX^56^ (Fig. 4c and Extended Data Fig. 12).

For the identification and reconstruction of the 100S disome, starting from the ribosome particle set that yielded the final 4.1 Å reconstruction, we inspected the subtomogram averages for evidence of higher-order ribosome assemblies. At low threshold levels, additional density was observed adjacent to a subset of ribosomes that was consistent with the presence of 100S disomes formed through binding of the long form of the hibernation-promoting factor (HPF). To identify candidate disome particles, custom Python scripts were used to shift ribosome particle coordinates to the putative position of the HPF interaction interface. For each ribosome particle, the distance to the nearest neighbouring ribosome was calculated using Euclidean distances between particle coordinates in tomogram space. Particles with a nearest-neighbour distance below 30 Å were classified as candidate disome-associated ribosomes. This procedure identified 4,114 ribosome particles.

These particles were mapped back into the tomograms and visually inspected to confirm that the corresponding ribosomes formed paired assemblies consistent with 100S disomes. The coordinates of the interacting ribosome partners were then recentered onto the putative disome interface and 2,057 disome-centered subtomograms were extracted from the refined tomograms in WarpTools at a pixel size of 3.82 Å.

Subtomograms were aligned using gold-standard 3D refinement in RELION 4.0.2^53^, initially without imposed symmetry and subsequently with C2 symmetry, resulting in a reconstruction with a global resolution of 9.9 Å. The refined particle parameters were subsequently imported into M and subjected to additional refinement cycles as described above for BCRII, including particle pose refinement, image warp correction, and CTF refinement. This yielded a final reconstruction of the 100S disome at 8.8 Å global resolution (Extended Data Fig. 11c).

To improve map interpretability, subtomogram averages were post-processed with LocSpiral^57^. Fitting of ETF in the local map of the ETF-BamD subcomplex (Fig. 4f) was performed by flexible fitting of an AlphaFold3^58^ prediction of a complex between the non-bifurcating ETF from *Geobacter metallireducens* GS-15 (GenBank ABB32381.1, ABB32382.1) and BamD using Isolde^39^.

### Tomographic volumes post-processing and visualization

For tomogram denoising and missing wedge reconstruction, the DeepDeWedge workflow^59^ was applied to M-refined half-tomograms, using subtomograms extracted individually from each reconstructed tomogram. Training was performed for 1000 epochs using subtomograms of size 80 × 80 × 80 pixels. Boundary masks were generated using Slabify^60^ from boundaries defined manually in IMOD^61^, delineating the lamella volume and excluding approximately 25–30 nm from both the upper and lower lamella surfaces. Subtomograms were extracted using strides of 80, 80 and 64 pixels along the X, Y and Z axes, respectively. The resulting tomograms were multiplied by the boundary mask and used as input for membrane segmentations in MemBrain-seg^62^. Storage granules were manually segmented in Amira (Thermo Fisher Scientific). For visualization, tomograms, segmentations and particles were imported and visualised in ArtiaX^56^. The segmentations were smoothed using 50 iterations of ‘*volume surfaceSmoothing’* in ChimeraX^63^.

### Estimation of intracellular particle concentrations

To estimate intracellular particle concentrations, cytoplasmic volumes were calculated from membrane segmentations generated using MemBrain-seg. These segmentations were used to define the cytoplasmic boundaries within each tomogram. A custom Python script was used to convert the number of voxels contained within the cytoplasmic segmentation into volumes expressed in liters. Particle coordinates from the final subtomogram averaging star files were used to determine the number of BCRII complexes and ribosomes located within the segmented cytoplasmic regions. The resulting intracellular particle concentrations across individual tomograms are summarised in Extended Data Fig. 11f,g.

### Synthesis of CoA esters

The synthesis of benzoyl-CoA, Ch1CoA, crotonyl-CoA, butyryl-CoA and glutaryl-CoA was achieved by coupling Li3CoA (20.0 mg, 24.4 µmol) with the respective carboxylic acids (51.2 µmol, 2.1 eq.) following the general procedure by Schachter and Taggart^64^. The free acids were activated with O-(Benzotriazol-1-yl)-N,N,N′,N′-tetramethyluronium tetrafluoroborate (TBTU, 17.2 mg, 53.7 µmol, 2.2 eq.), 1-hydroxybenzotriazole (HOBt, 7.2 mg, 53.7 µmol, 2.2 eq.) and N,N-Diisopropylethylamine (DIPEA, 21.3 mg, 28.7 µL, 165 µmol, 6.8 eq.) in DMF (1.8 mL) at room temperature under constant stirring for 30 min. Then, 200 µL Li_3_CoA in 100 mM NaHCO_3_ (pH 8.2) was added and the mixture was incubated for 30 min at room temperature under constant stirring. After dilution to 10 mL in 10 mM K_2_HPO_4_ (pH 6.8), the CoA ester was purified by reversed-phase high performance liquid chromatography (HPLC) on a Waters 1525 preparative HPLC using a Waters, Atlantis® T3 Prep OBD™, column (5 µm, 19 × 250 mm). Compounds eluted in a linear acetonitrile (ACN)/10 mM potassium phosphate (pH 6.8) between 2-42 % ACN (8 mL min^−1^). Purity of elution fractions was analysed by UPLC analysis. Selected fractions were frozen in liquid nitrogen and lyophilised^65^. The synthesis of 1,5-dienoyl-CoA was based on the enzymatic conversion of previously synthesised Ch1CoA (50 µmol), using cyclohexenoyl-CoA DH from *G. metallireducens* (Gmet_3306, 0.5-1.5 mg mL^−1^) as the catalyst and 2 mM potassium hexacyanoferrate(III) as the electron acceptor in 100 mM MOPS/KOH (pH 7.5, 20 mM total volume) under anoxic conditions. The reaction was incubated for 15 min at 30 °C, stopped by the addition of 40 mL methanol, and subsequently purified by preparative HPLC as described above.

### Phylogenetic consensus tree of BamDs

To create the phylogenetic tree of BamDs, a multiple sequence alignment of BamDs identified by Anselmann^17^ was performed using Clustal Omega via the EMBL-EBI framework^66^. The phylogenetic tree was created by MEGA12^67^. The Maximum Likelihood method and Jones-Taylor-Thornton (JTT) matrix-based model^68^ were employed with a bootstrap^69^ value of 1,000. The initial tree for the heuristic search was selected by choosing the tree with the superior log-likelihood between a Neighbor-Joining (NJ)^70^ and Maximum Parsimony (MP) tree. The NJ tree was generated using a matrix of pairwise distances computed using the JTT model. The MP tree had the shortest length among 10 independent MP tree searches. The analytic procedure encompassed 39 amino acid sequences with 475 positions in the final dataset.

## Acknowledgements

This research was supported by Deutsche Forschungsgemeinschaft (SFB 1381, project ID 403222702 to L.A. and M.B.). We thank Sebastian Estelmann for providing the plasmids encoding the genes for ETF and BDH. J.M.S. acknowledges the Deutsche Forschungsgemeinschaft for an Emmy Noether grant (SCHU 3364/1-1), RTG 2937 and the European Union’s Horizon 2020 research and innovation programme (Two-CO2-One; grant agreement no. 101075992). The views and opinions expressed are those of the author(s) only and do not necessarily reflect those of the European Union or the European Research Council. Neither the European Union nor the granting authority can be held responsible for them. D.T. and T.C.P were supported by EMBO Long Term Fellowships. We thank W. Wietrzynski and S. Zufferey for their support with grid preparation for cryo-ET. We thank R.D. Righetto for support during cryo-ET data processing. Cryo-ET instrumentation and computational resources were provided by the University of Basel BioEM Lab and sciCORE, respectively. This paper was typeset with the bioRxiv word template by @Chrelli: www.github.com/chrelli/bioRxiv-word-template

## Author contributions

M.B. conceptualised the project. M.B., B.D.E., J.M.S. and U.E. coordinated the experiments. L.A. cultured cells, expressed and purified the proteins, designed, carried out and analysed the enzyme assays, synthesised CoA esters and constructed the phylogenetic tree. T.K. prepared the plasmid encoding BamGHI and performed preliminary heterologous expression experiments. A.K. optimised the sample conditions for cryo-EM SPA. A.K. and T.R.T. prepared cryo-EM grids. K.K. collected and processed initial screening dataset. A.K. and S.B. acquired the high-resolution cryo-EM dataset. A.K. processed the cryo-EM dataset, performed local refinements, built and refined the final models. A.K. analysed and interpreted the cryo-EM models. L.A., A.K. and T.C.P. prepared grids for cryo-ET. D.T. performed FIB milling and cryo-ET data acquisition. D.T. and T.C.P. processed cryo-ET data, carried out subtomogram averaging, cryo-ET data analysis and figures. M.B., B.D.E., J.M.S., U.E., L.A., A.K., D.T. and T.C.P. wrote the manuscript together with insights from all other authors. L.A., A.K., D.T. and T.C.P. contributed equally to the manuscript. M.B., B.D.E. and J.M.S. are the corresponding authors.

## Data availability

The cryo-EM maps reported in this article have been deposited in the Electron Microscopy Data Bank under accession code EMD-57300 (BCRII complex full assembly - composite map), EMD-57318 (BCRII complex state 1 - composite map), EMD-57334 (BCRII complex state 2 - composite map), EMD-57349 (BamED protomer local refined map), EMD-57350 (BamED dimer local refined map), EMD-57351 (BamD dimer local refined map), EMD-57352 (BamBC local refined map), EMD-57353 (BamGHIF Arm 1 State 1 local refined map), EMD-57354 (BamED Arm 1 State 1 local refined map), EMD-57355 (BamGHIF Arm 1 State 2 local refined map), EMD- 57356 (BamED Arm 1 State 2 local refined map), EMD-57357 (BamGHIF Arm 2 local refined map). The atomic models have been deposited in the Protein Data Bank (PDB) under the codes 29QY (BCRII complex full assembly - composite model), 29RC (BCRII complex state 1 - composite model), and 29RN (BCRII complex state 2 - composite model). Cryo-ET cellular tomograms are available in the Electron Microscopy Data Bank (EMDB) with the accession codes EMD-XXXXT1-XXXT8. Raw electron tomography data are available in the Electron Microscopy Public Image Archive (EMPIAR-XXXXX). Cryo-ET structures have been deposited to the Electron Microscopy Data Bank under accession numbers EMD-XXXS1 (class II BCR complex), EMD-XXXS2 (ETF-BamD), EMD-XXXS3 (ribosome), EMD-XXXS4 (100S disome). The source data underlying the main text, Figs. 2 and 4, and Extended Data Figs. 2, 9 and 13 are provided in the Supplementary Data files. The mass spectrometry protein identification data have been deposited to the ProteomeXchange Consortium via the PRIDE^71^ partner repository with the dataset identifier PXD076380. Additional data that support the findings of this study are available from the corresponding authors upon request. Supplementary Data files are provided with this paper.

