## Extended Figures 1-13, Extended Table 1-2 for "Coupling of electron-bifurcation modules powers aromatic ring reduction beyond the biological redox window"

---

**Extended Data Figures 1-13**

**Extended Data Tables 1-2**

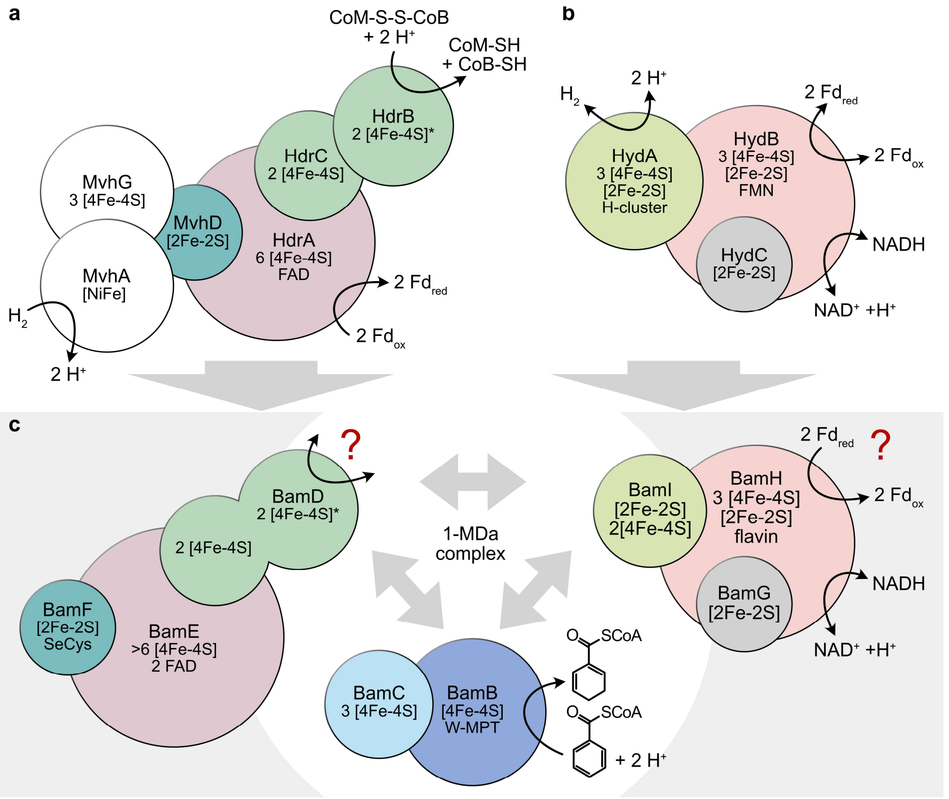

**Extended Data Fig. 1 | Schematic representation of the modular architecture, cofactors and proposed FBEB processes of (a) electron-bifurcating heterodisulfide reductase–[NiFe]-hydrogenases, (b) electron-confurcating/bifurcating hydrogenases, and (c) class II benzoyl-CoA reductases (BCRII).** Homologous subunits are shown in identical colours. The connectivity of the BamDEF, BamBC and BamGHI modules in BCRII remains unresolved. Left panel: similarities between BCRII and FBEB modules of heterodisulfide reductases. BamE is homologous to the electron-bifurcating HdrA subunit; BamF resembles MvhD and likely functions as an adaptor to an unidentified electron-input module; BamD can be considered a fusion of the electron-transferring and catalytic HdrB/HdrC subunits. Whether BamD interacts with an external electron acceptor/donor is unknown. Right panel: similarities between BCRII and FBEB modules of hydrogenases. BamH is homologous to the electron-bifurcating/confurcating HydB subunit; BamG and BamI correspond to the electron-transferring HydA/HydC subunits but lack the H-cluster-binding domain. The catalytic BamB/BamC subunits belong to the tungsten-cofactor-containing aldehyde:ferredoxin oxidoreductase family.

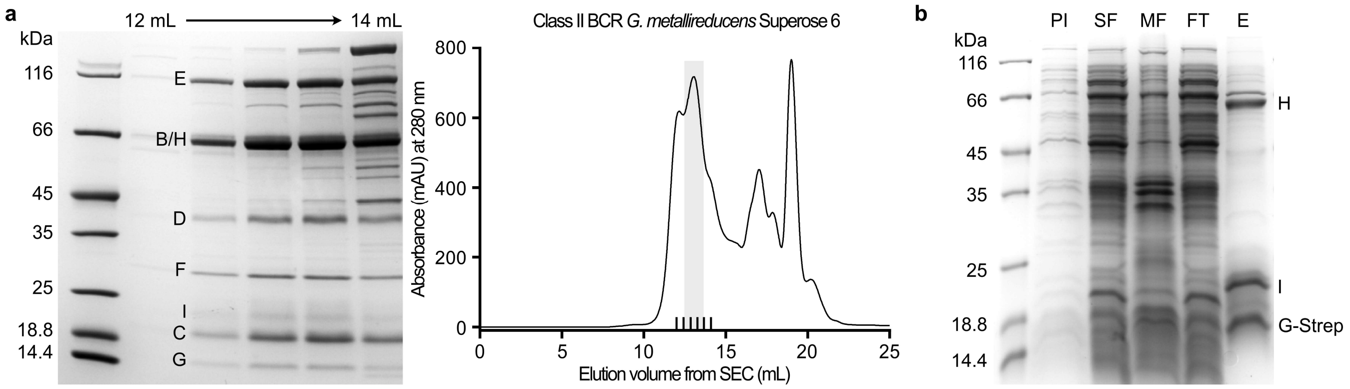

**c**

| Cofactor content (mol mol-1) | | |
|---|---|---|
| | Class II BCR complex | BamGHI |
| FAD | 4.6 ± 1.0 | - |
| FMN | 0.75 ± 0.36 | 0.18 ± 0.16 |
| Fe | 183 ± 33 | 25.9 ± 7.2 |

**d**

| Specific activities (µmol mL$^{-1}$mol$^{-1}$) | | |
|---|---|---|
| | Class II BCR complex | BamGHI |
| DCO | 20.1 ± 4.6 | - |
| NOR | 0.49 ± 0.09 | 0.046 |
| MV:NAD$^+$ | 1.59 ± 0.54 | - |
| Fd:NAD$^+$ | 0.0 | - |
| DCoA:NAD$^+$ | 0.0 | - |

**e**

| | Gene identifier | Protein identifier |
|---|---|---|
| BamB-1* | Gmet_2087 | Q39TV8 |
| BamB-2 | Gmet_1082 | Q39UP3 |
| BamC-1* | Gmet_2086 | Q39TV9 |
| BamC-2 | Gmet_1803 | Q39UP2 |
| BamD | Gmet_2085 | Q39TW0 |
| BamE | Gmet_2084 | Q39TW1 |
| BamF | Gmet_2083 | Q39TW2 |
| BamG | Gmet_2081 | Q39TW4 |
| BamH | Gmet_2080 | Q39TW5 |
| BamI | Gmet_2079 | Q39TW6 |

**Extended Data Fig. 2 | Enrichment of BCRII and BamGHI from *G. metallireducens*, including subunit identifiers, cofactor content and enzymatic activities. a,** SDS-PAGE (left) and size-exclusion chromatography (SEC, right) profile (Superose 6 Increase 10/300 GL, 24 mL) of the *G. metallireducens* BCRII complex purified from the 350 mM KCl fraction during DEAE Sepharose chromatography. **b,** SDS-PAGE of BamGHI from *G. metallireducens* after Strep-tag affinity purification. Samples: pre-induction cells (PI), soluble fraction (SF), membrane fraction (MF), flow-through (FT) and eluate (E). **c,** Quantification of FAD, FMN and Fe content. **d,** Specific activities of the enriched complexes. DCO = 1,5-dienoyl-CoA:methyl viologen (MV) oxidoreductase, NOR = NADH:benzyl viologen oxidoreductase, MV:NAD$^+$ = $MV_{red}$:NAD$^+$ oxidoreductase. **e,** Subunit identifiers of BCRII from *G. metallireducens*. From the two gene copies encoding BamBC variants, the gene products marked with * were those detected, together with the BamDEFGHI gene products, by mass spectrometry analysis in the SDS-gel shown in **a**.

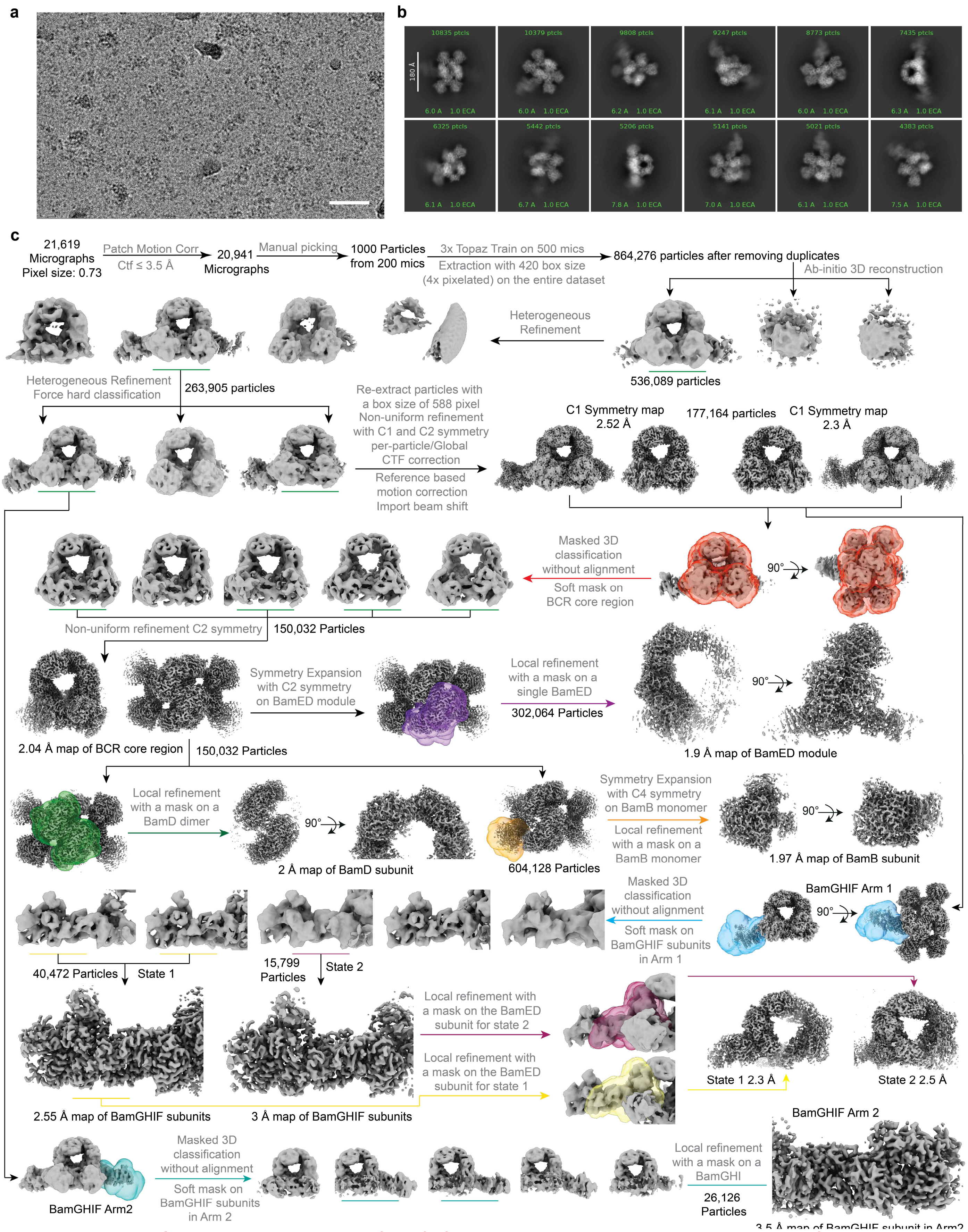

**Extended Data Fig. 3 | Cryo-EM data processing workflow of BCRII complex incubated with NADH and pre-reduced ferredoxin. a,** A representative motion-corrected cryo-EM micrograph from the full dataset comprising 21,619 images (scale bar = 50 nm). The micrographs show BCRII complexes vitrified under strictly anaerobic conditions in presence of NADH and pre-reduced ferredoxin. **b,** Reference-free 2D class average revealing the characteristic omega shaped core of the BCRII complex flanked by flexible peripheral arms in multiple orientations. **c,** Data-processing scheme used to obtain high-resolution reconstructions of the core assembly and its associated arms, including classification of particles into distinct conformational states.

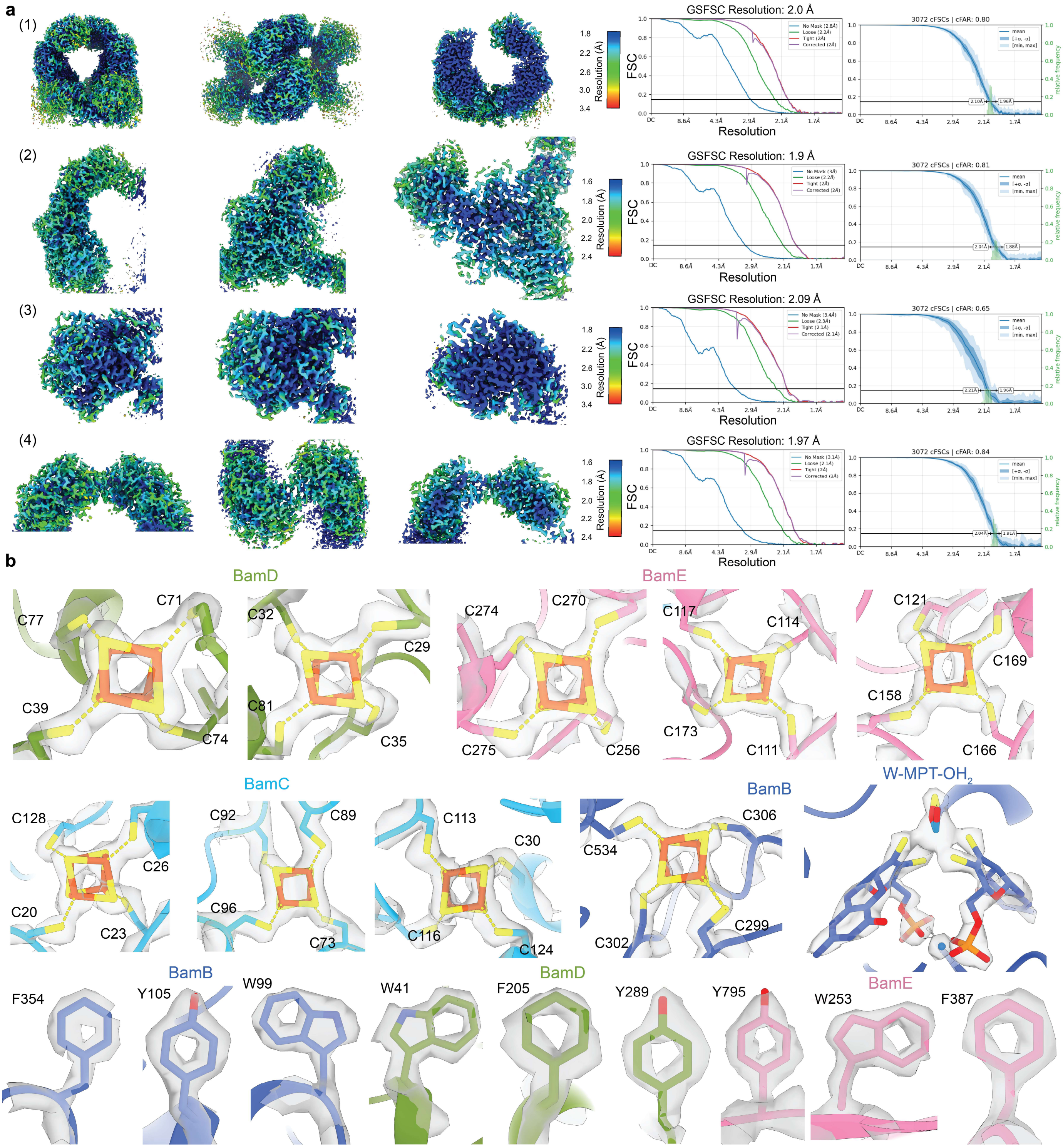

**Extended Data Fig. 4 | Cryo-EM maps resolution, orientation diagnostics, and model-to-map fit for the BamFEDBC region. a,** Left panel: Cryo-EM maps coloured according to local resolution as calculated in CryoSPARC. Two orthogonal views (left and middle) together with a cutaway view of the central region (right) are shown for: (1) dimeric core of the BCRII complex, (2) BamED subunit, (3) BamB subunit, and (4) dimeric BamD subunits. Right panel: Plots showing resolution estimates using gold-standard Fourier shell correlation (GSFSC) together with orientation diagnostics derived from canonical FSC analysis. **b,** Cryo-EM map surrounding cofactors, Fe−S clusters, and certain aromatic residues present within the core region of the BCRII complex, highlighting the quality of reconstructions.

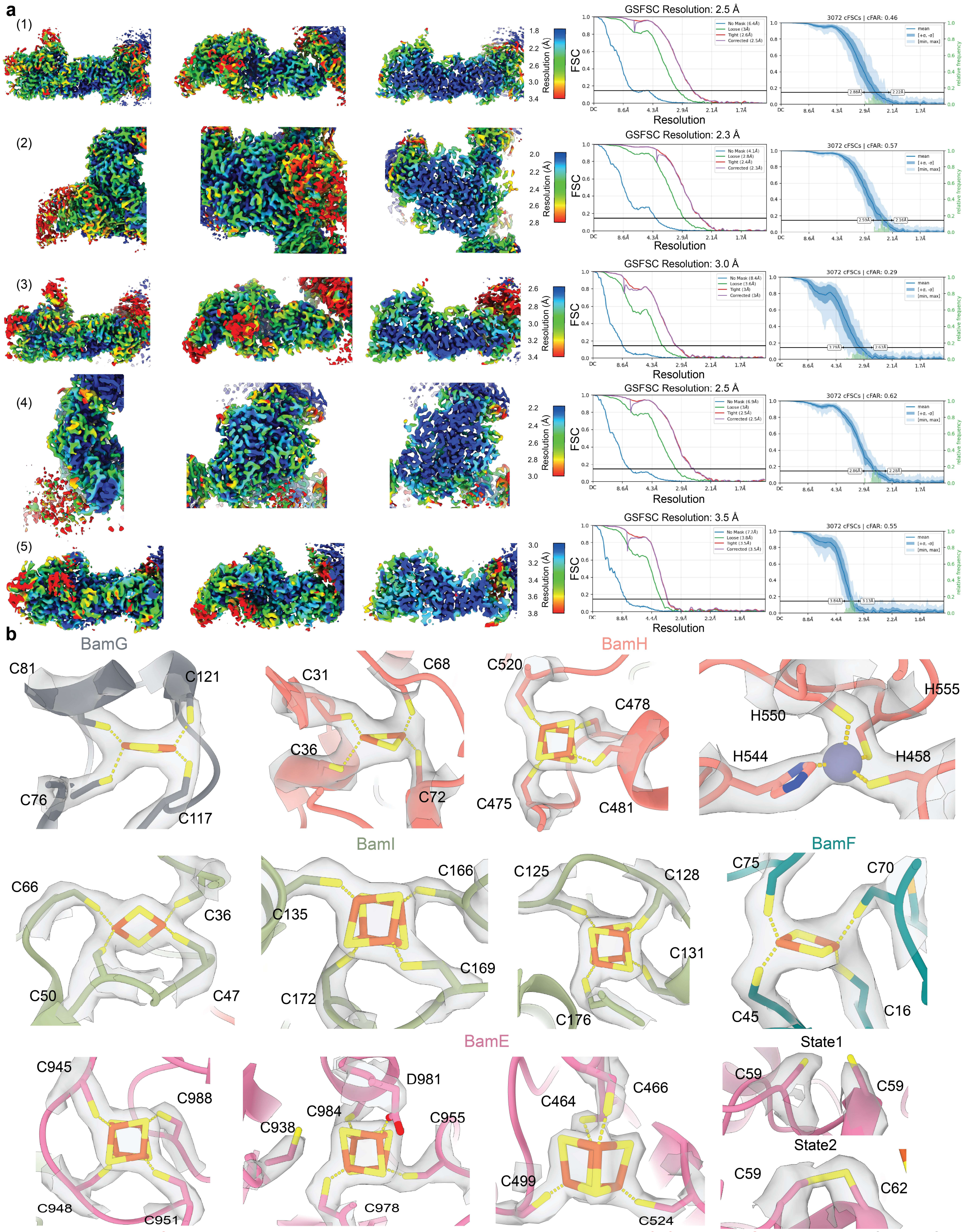

**Extended Data Fig. 5 | Cryo-EM maps resolution, orientation diagnostics, and model-to-map fit for the BamGHIF region. a,** Left panel: Cryo-EM maps coloured according to local resolution as calculated in CryoSPARC. Two orthogonal views (left and middle) together with a cutaway view of the central region (right) are shown for: (1) BamGHIF arm 1 state 2, (2) BamEF subunit for state 2, (3) BamGHIF arm 1 state 1, (4) BamEF subunit for state 1, and (5) BamGHIF arm 2 (second arm located in the right side of the complex). Right panel: Plot showing resolution estimates using gold-standard Fourier shell correlation (GSFSC) together with orientation diagnostics derived from canonical FSC analysis. **b,** Cryo-EM map surrounding Fe−S clusters present within the BamGHIF region of the BCRII complex.

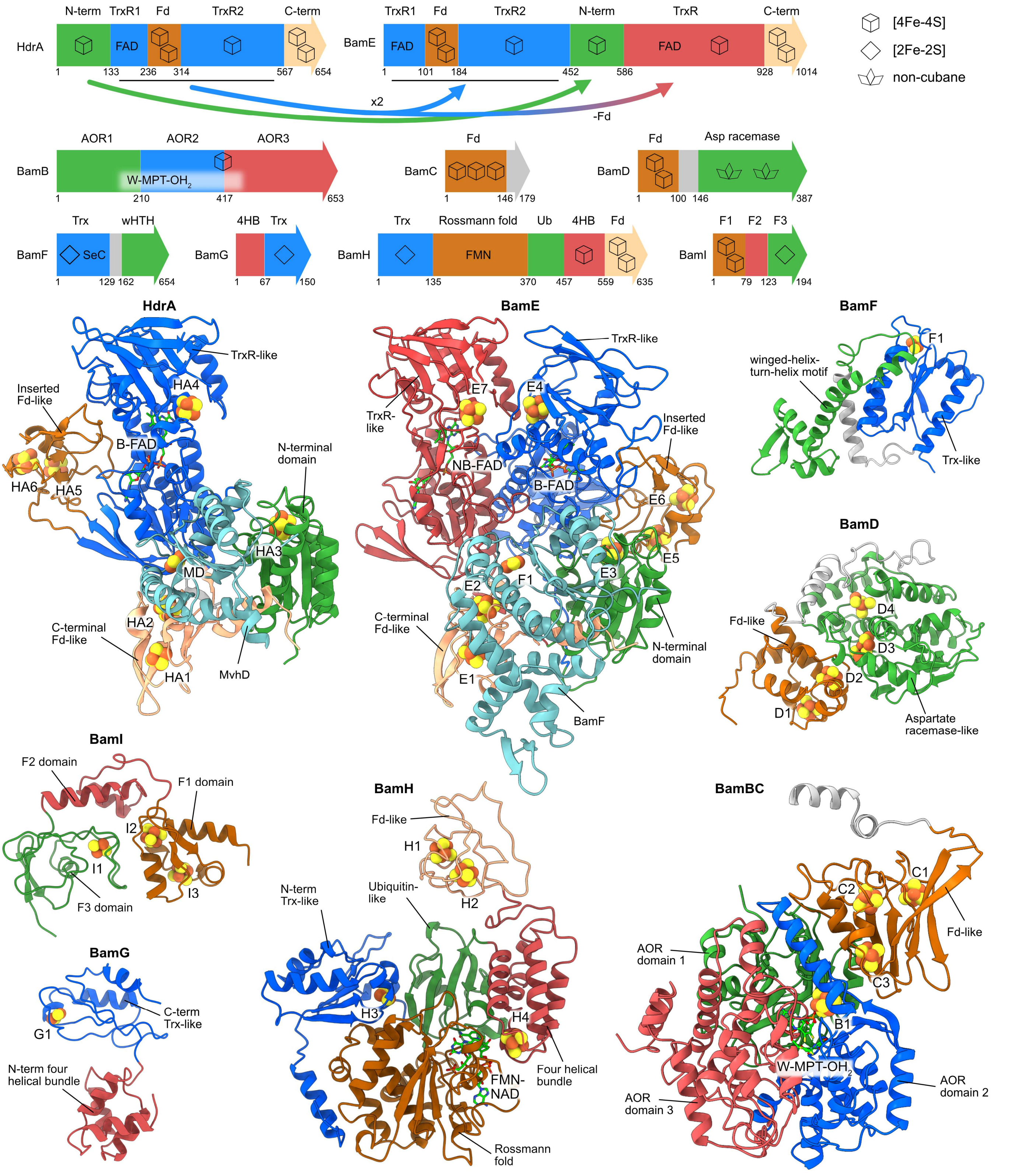

**Extended Data Fig. 6 | Domain dissection of the BCRII complex.** Domain distribution is coloured according to the respective structural folds. HdrA from *Methanothermococcus thermolithotrophicus* is compared to BamE, to illustrate the domain rearrangements between the two subunits. 4HB, four helical bundle domain; AOR1-3, aldehyde:ferredoxin oxidoreductase-like domain 1-3; Asp racemase, aspartate racemase-like domain, C-ter, C-terminal domain; F1-3, [FeFe] hydrogenase-like domain 1-3; Fd, ferredoxin-like domain; N-ter, N-terminal domain; Trx, thioredoxin-like domain; TrxR, thioredoxin reductase-like domain; wHTH, winged-helix-turn-helix motif.

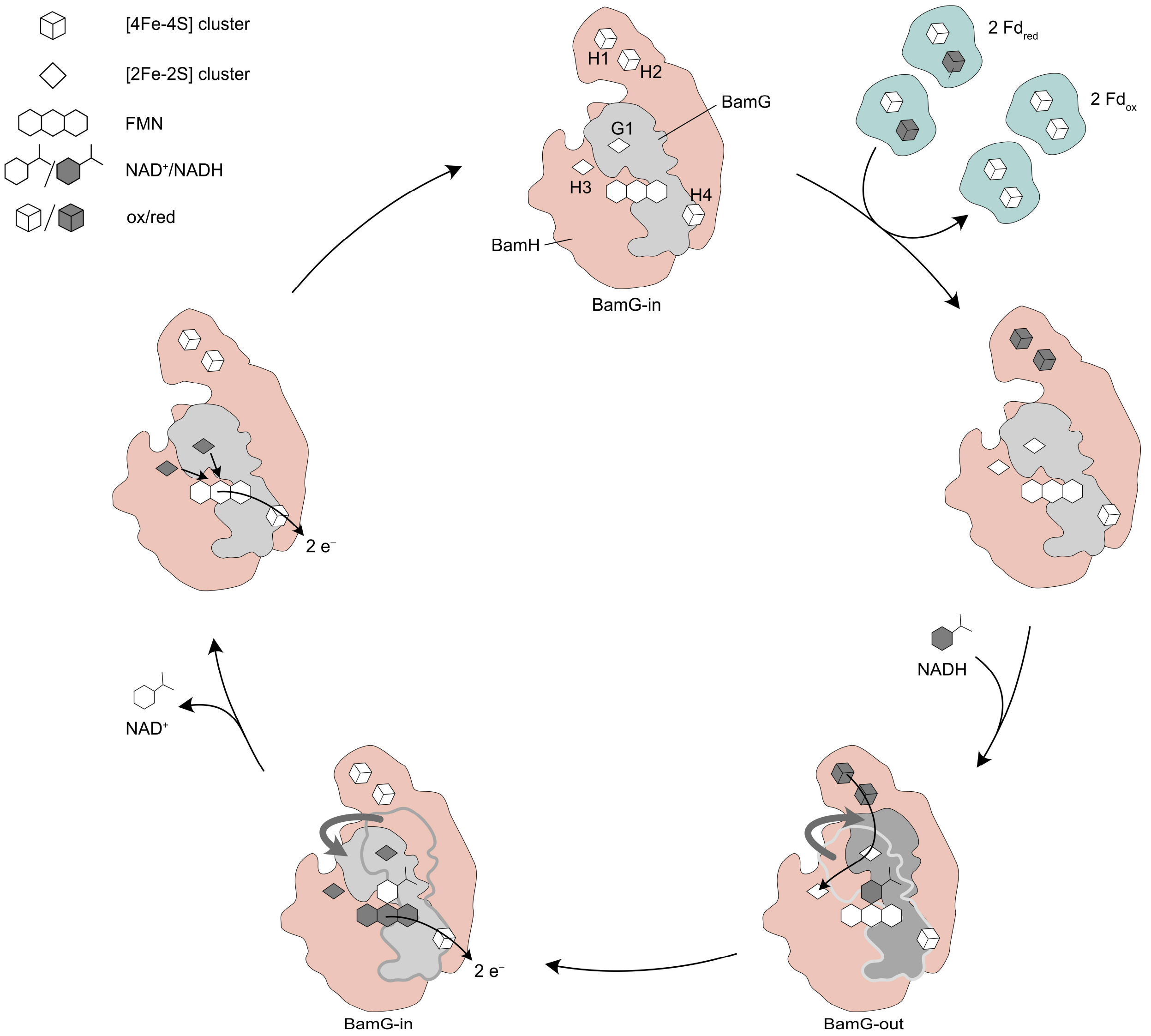

**Extended Data Fig. 7 | Model for the electron confurcation mechanism in BamGH.** The reaction is initiated by $Fd_{red}$-dependent reduction of clusters H1/H2. In this BamG-in (closed) state, electron transfer from cluster H2 to G1 cluster is blocked. The binding of NADH triggers a conformational transition to the BamG-out (open) state, enabling the reduction of clusters G1/H3 by reduced H1/H2 clusters. After electron transfer, BamG relaxes back into the BamG-in state. The electrons derived from NADH are transferred towards H4 and the BamI Fe–S clusters accompanied by $NAD^+$ dissociation. Next, electrons residing on G1/H3 are transferred toward BamI via FMN and cluster H4. Alternative order of events during the rearrangement from the BamG-out to the BamG-in state are conceivable[22,30].

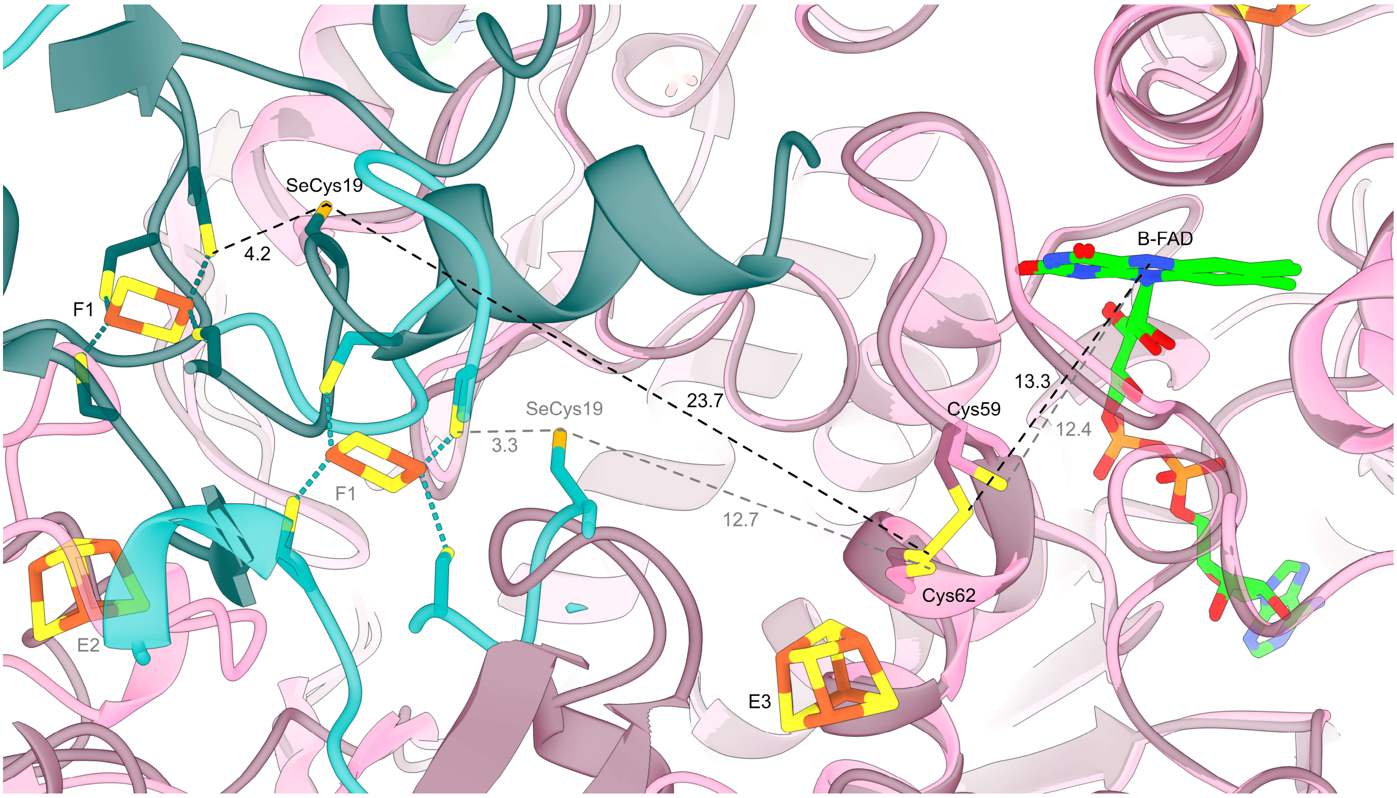

**Extended Data Fig. 8 | Enlarged view of spatial arrangement and distances between cofactors in states 1 and 2 of the BamEF region.** BamF and BamE are shown in lighter (state 1) and darker (state 2) shades of blue and pink, respectively. Labels and distances are shown in grey for state 1 and black for state 2. All the distances are given in Angstrom (Å).

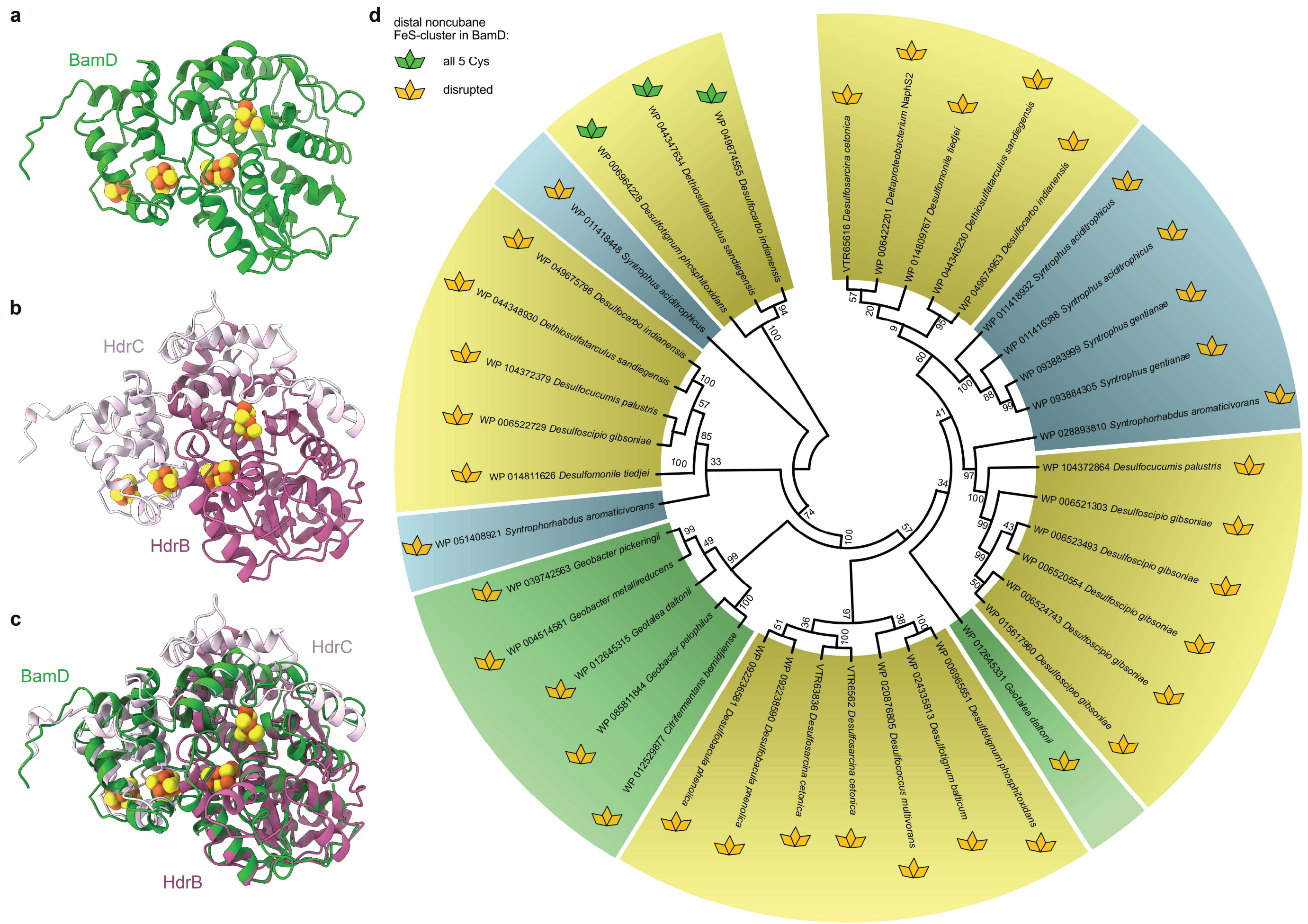

**Extended Data Fig. 9 | Comparison of BamD with the HdrBC subcomplex. a,** Atomic model of BamD, and, **b,** HdrBC from *Methanothermococcus thermolithotrophicus* (PDB 5ODC). **c,** Structural superposition of BamD with HdrBC (RMSD 1.031 Å). **d,** Phylogenetic tree of BamDs from BCRII-containing organisms. BamDs from metal-reducers are shown in green, sulfate-reducing organisms in yellow, and syntrophic organisms in blue. Conserved amino acid residues coordinating the D4/HB2 cluster are indicated by symbols coloured green (all cysteine residues conserved) or yellow (≥ 1 cysteine residue absent). The phylogenetic tree was based on a multiple sequence alignment (MSA) produced with Clustal Omega[66] and constructed in MEGA12[67] as described in the materials and methods section.

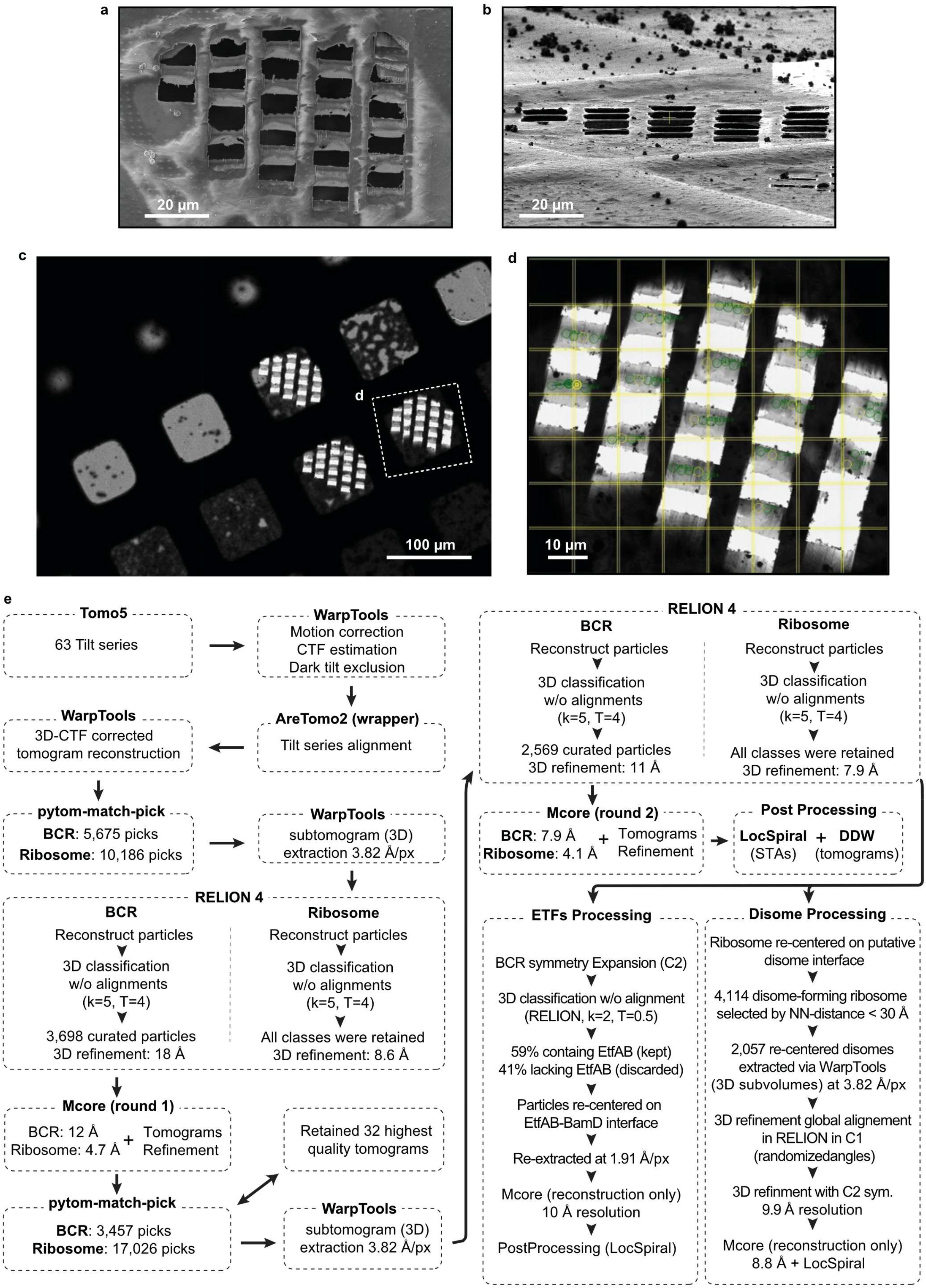

a
20 µm
b
20 µm
c
d
100 µm
d
10 µm
e
Tomo5
63 Tilt series
WarpTools
Motion correction
CTF estimation
Dark tilt exclusion
AreTomo2 (wrapper)
Tilt series alignment
WarpTools
3D-CTF corrected
tomogram reconstruction
pytom-match-pick
BCR: 5,675 picks
Ribosome: 10,186 picks
WarpTools
subtomogram (3D)
extraction 3.82 Å/px
RELION 4
BCR
Reconstruct particles
3D classification
w/o alignments
(k=5, T=4)
3,698 curated particles
3D refinement: 18 Å
Ribosome
Reconstruct particles
3D classification
w/o alignments
(k=5, T=4)
All classes were retained
3D refinement: 8.6 Å
Mcore (round 1)
BCR: 12 Å
Ribosome: 4.7 Å
+
Tomograms
Refinement
Retained 32 highest
quality tomograms
pytom-match-pick
BCR: 3,457 picks
Ribosome: 17,026 picks
WarpTools
subtomogram (3D)
extraction 3.82 Å/px
RELION 4
BCR
Reconstruct particles
3D classification
w/o alignments
(k=5, T=4)
2,569 curated particles
3D refinement: 11 Å
Ribosome
Reconstruct particles
3D classification
w/o alignments
(k=5, T=4)
All classes were retained
3D refinement: 7.9 Å
Mcore (round 2)
BCR: 7.9 Å
Ribosome: 4.1 Å
+
Tomograms
Refinement
Post Processing
LocSpiral + DDW
(STAs) (tomograms)
ETFs Processing
BCR symmetry Expansion (C2)
3D classification w/o alignment
(RELION, k=2, T=0.5)
59% containg EtfAB (kept)
41% lacking EtfAB (discarded)
Particles re-centered on
EtfAB-BamD interface
Re-extracted at 1.91 Å/px
Mcore (reconstruction only)
10 Å resolution
PostProcessing (LocSpiral)
Disome Processing
Ribosome re-centered on putative
disome interface
4,114 disome-forming ribosome
selected by NN-distance < 30 Å
2,057 re-centered disomes
extracted via WarpTools
(3D subvolumes) at 3.82 Å/px
3D refinement global alignement
in RELION in C1
(randomizedangles)
3D refinment with C2 sym.
9.9 Å resolution
Mcore (reconstruction only)
8.8 Å + LocSpiral

**Extended Data Fig. 10 | Cryo-ET data acquisition and processing pipeline. a,** Scanning electron microscopy (SEM) image of a grid square after cryo-focused ion beam (cryo-FIB) milling. **b,** Ion beam image of the same lamellae. **c,** Low-magnification cryo-TEM overview of a grid containing multiple lamellae. **d,** Positions for automated tilt-series acquisition across multiple lamellae within a single grid square, defined in Tomo5 software. **e,** Cryo-ET data processing flowchart used for *in situ* subtomogram averaging, tomogram refinement, and post-processing.

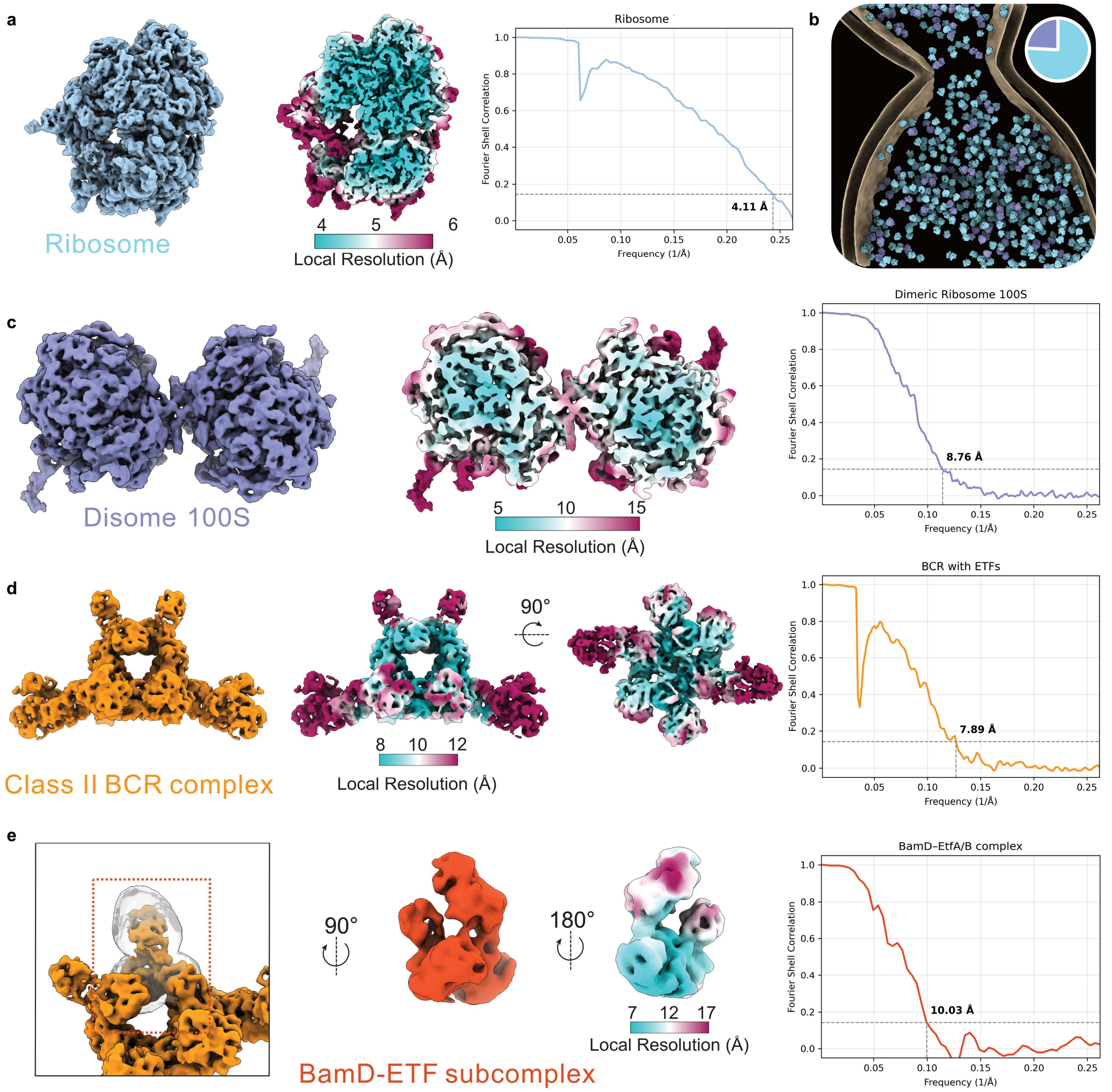

f

| | BCR (particles) | Ribosome (particles) | Imaged cytoplasm (fL) | Ribosome (μM) | BCR (μM) |
|---|---|---|---|---|---|
| Tomo 01 | 145 | 793 | 0.089 | 14.790 | 2.704 |
| Tomo 02 | 95 | 963 | 0.069 | 23.150 | 2.284 |
| Tomo 03 | 190 | 715 | 0.080 | 14.785 | 3.929 |
| Tomo 04 | 145 | 659 | 0.048 | 22.635 | 4.980 |
| Tomo 05 | 193 | 760 | 0.077 | 16.488 | 4.187 |
| Tomo 06 | 126 | 757 | 0.093 | 13.555 | 2.256 |
| Tomo 07 | 65 | 537 | 0.072 | 12.384 | 1.499 |

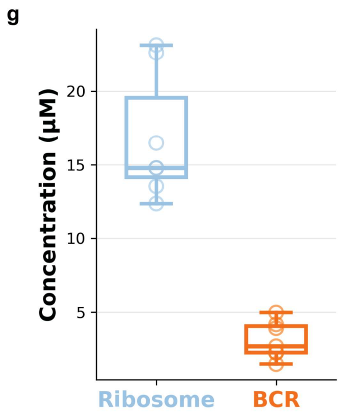

**Extended Data Fig. 11 | Subtomogram averaging map quality and intracellular particle concentrations. a,** Ribosome subtomogram average, sliced-through map coloured by local resolution, and masked Fourier shell correlation (FSC) curve. **b,** Back-projections of ribosome (cyan) and disome (purple) particles within a representative tomogram, with the overall proportions of each species in the dataset indicated. **c,** 100S disome subtomogram average, sliced-through map coloured by local resolution, and masked FSC curve. **d,** BCRII subtomogram average, map coloured by local resolution, and masked FSC curve. **e,** Mask defining the ETF-BamD subcomplex, corresponding focused map reconstructed from particles with occupied ETF-binding sites, map coloured by local resolution, and masked FSC curve. **f,** Particle concentration analysis performed on seven representative high-quality tomograms. **g,** Box plot showing the estimated intracellular concentrations of BCRII and ribosomes across the seven tomograms analysed in **f**. Points indicate individual tomograms. The reported resolutions correspond to the FSC 0.143 criterion.

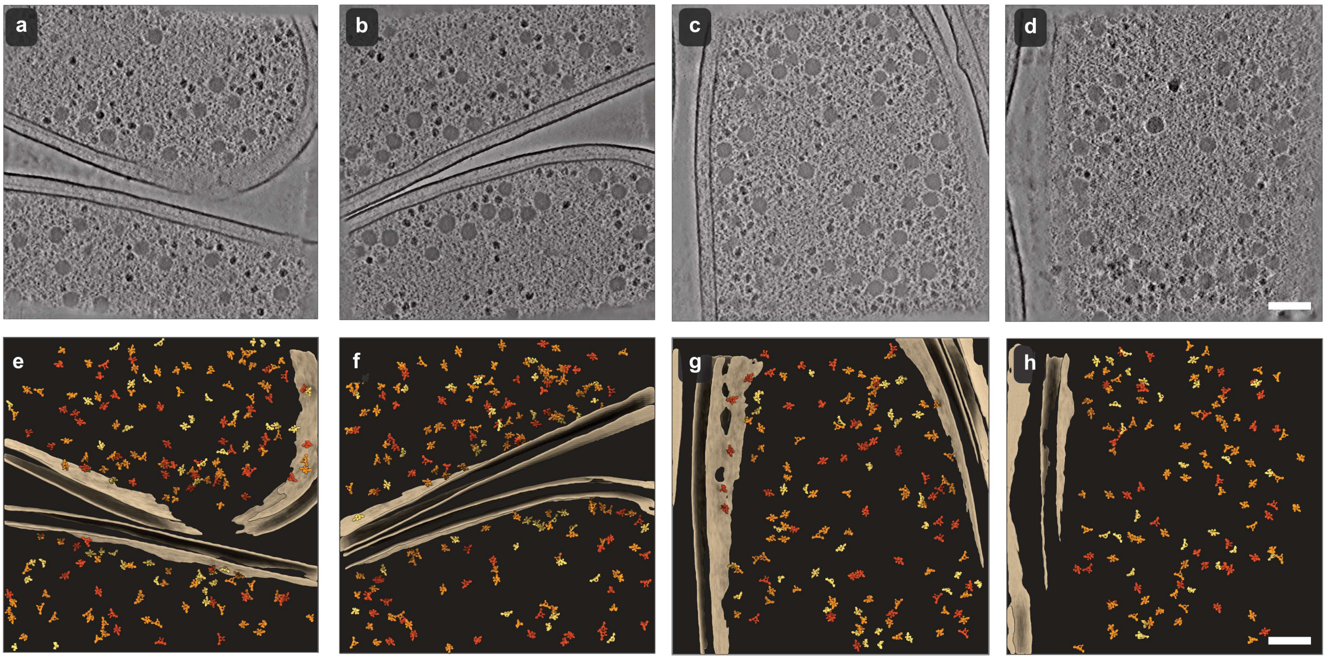

**Extended Data Fig. 12 | Spatial distribution of ETF occupancy states.** Four representative tomograms and the corresponding back-projections of BCRII particles. Colours indicate ETF occupancy at each BCRII complex: yellow is 0× ETF; orange is 1× ETF; red is 2× ETF. Membranes in tan. Scale bar, 100 nm.

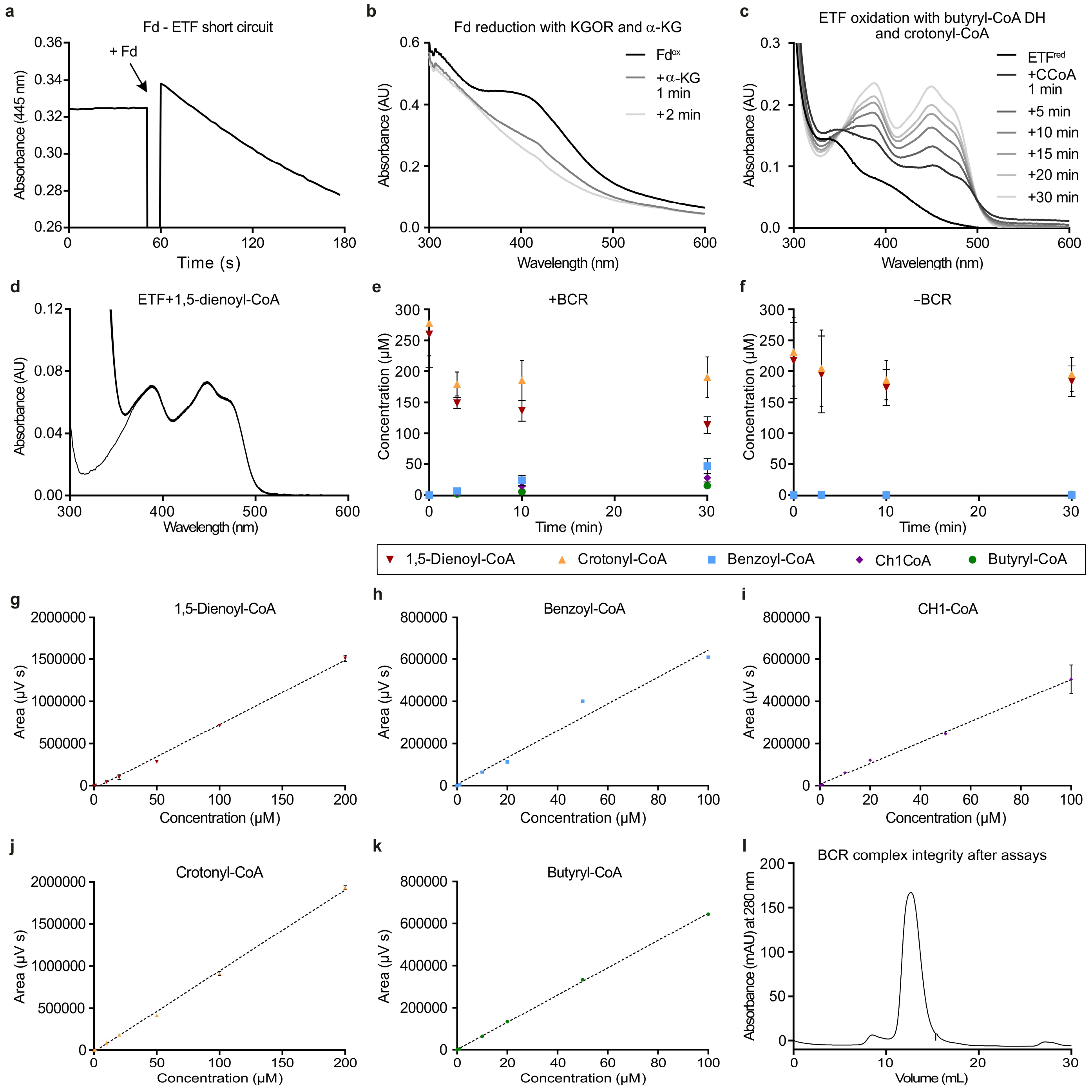

**Extended Data Fig. 13 | Short circuit and regeneration systems in electron confurcation/bifurcation assays with BCRII, as well as electron transfer between BCRII and ETF. a,** Short circuit reaction between reduced Fd (1 µM) and ETF (20 µM), and **b,** Reduction of Fd (14 µM) by KGOR (1.7 µM) in the presence of CoA (1 mM) and α-ketoglutarate (1 mM). **c,** Oxidation of pre-reduced ETF (20 µM) with butyryl-CoA dehydrogenase (1 µM) from *Desulfosarcina cetonica* and crotonyl-CoA (100 µM). **d,** UV/vis spectra of the control assay, demonstrating that ETF (10 µM) is not reduced by 1,5-dienoyl-CoA (150 µM) in the absence of BCRII. **e,** Discontinuous *in vitro* assays coupling ETF reduction by BCRII, using 1,5-dienoyl-CoA as electron donor, to the reduction of crotonyl-CoA to butyryl-CoA with ETF by butyryl-CoA dehydrogenase. Samples were collected after 0, 3, 10 and 30 min, quenched in methanol and analysed by UPLC. **f,** Control assays performed in the absence of BCRII. **g-k,** Calibration curves for CoA esters prepared and analysed identically to assay samples. Assays data are mean ± s.d. of three replicates. Calibration curves were measured in two technical replicates and shown as mean ± s.d. **l,** Superose 6 size-exclusion chromatography demonstrating the integrity of the BCR complex 24 h after enrichment when stored at 10 °C. Abbreviations: 1,5-dienoyl-CoA, cyclohexa-1,5-diene-1-carbonyl-CoA; CH1-CoA, cyclohexa-1-ene-1-carbonyl-CoA.

**Extended Data Table 1 | Cryo-EM data collection, refinement, and validation statistics.**

| | **BCRII complex full assembly (composite map and model)** (EMDB - 57300) (PDB - 29QY) | **BCRII complex State-1 (composite map and model)** (EMDB - 57318) (PDB - 29RC) | **BCRII complex State-2 (composite map and model)** (EMDB - 57334) (PDB - 29RN) | **BamED protomer local refine** (EMDB - 57349) | **BamED dimer local refine** (EMDB - 57350) |
|---|---|---|---|---|---|
| **Data collection and processing** | | | | | |
| Magnification | 165,000 x | 165,000 x | 165,000 x | 165,000 x | 165,000 x |
| Voltage (kV) | 300 | 300 | 300 | 300 | 300 |
| Electron exposure (e–/Å$^2$) | 60.0 | 60.0 | 60.0 | 60.0 | 60.0 |
| Defocus range (μm) | 0.8-1.8 | 0.8-1.8 | 0.8-1.8 | 0.8-1.8 | 0.8-1.8 |
| Pixel size (Å) | 0.73 | 0.73 | 0.73 | 0.73 | 0.73 |
| Symmetry imposed | C1 | C1 | C1 | C1 | C1 |
| Initial particle images (no.) | 864,276 | 864,276 | 864,276 | 864,276 | 864,276 |
| Final particle images (no.) | 177,164 | 40,472 | 15,799 | 302,064 | 150,032 |
| Map resolution (Å) | | | | 1.9 | 2.0 |
| FSC threshold | | | | 0.143 | 0.143 |
| Map resolution range (Å) | | | | 1.8-2.0 | 1.96-2.1 |
| **Refinement** | | | | | |
| Initial model used (PDB code) | *de novo*, AlphaFold3 | *de novo*, AlphaFold3 | *de novo*, AlphaFold3 | - | - |
| Model resolution (Å) | | | | | |
| FSC threshold | | | | | |
| Model resolution range (Å) | | | | | |
| Map sharpening *B* factor (Å$^2$) | | | | 39.4 | 41.2 |
| Model composition | | | | | |
| Non-hydrogen atoms | 67386 | 20576 | 20576 | | |
| Protein residues | 8536 | 2627 | 2627 | - | - |
| Ligands | FMN: 2, NADH: 2, SEC: 2, FES: 8, SF4: 43, Zn: 2, FAD:4, 9S8: 2, MG:4, T7R: 4, 3S4: 2 | FMN: 1, NADH: 1, SEC: 1, FES: 4, SF4: 14, Zn: 1, FAD:2, 9S8: 1, 3S4: 1 | FMN: 1, NADH: 1, SEC: 1, FES: 4, SF4: 14, Zn: 1, FAD:2, 9S8: 1, 3S4: 1 | | |
| *B* factors (Å$^2$) | | | | - | - |
| Protein | 40.16 | 45.82 | 75.22 | - | - |
| Ligand | 30.62 | 38.92 | 66.89 | - | - |
| R.m.s. deviations | | | | - | - |
| Bond lengths (Å) | 0.006 | 0.005 | 0.015 | - | - |
| Bond angles (°) | 1.129 | 1.116 | 1.283 | - | - |
| Validation | | | | - | - |
| MolProbity score | 1.63 | 1.87 | 2.08 | - | - |
| Clashscore | 3.54 | 3.94 | 5.4 | - | - |
| Poor rotamers (%) | 1.71 | 2.14 | 2.3 | - | - |
| Ramachandran plot | | | | - | - |
| Favored (%) | 95.55 | 93.22 | 92.54 | - | - |
| Allowed (%) | 4.06 | 6.20 | 7.15 | - | - |
| Disallowed (%) | 0.39 | 0.57 | 0.31 | - | - |

| | BamD dimer local refine (EMDB - 57351) | BamBC local refine (EMDB - 57352) | BamGHIF Arm 1 State 1 local refine (EMDB - 57353) | BamED Arm 1 State 1 local refine (EMDB - 57354) | BamGHIF Arm 1 State 2 local refine (EMDB - 57355) | BamED Arm 1 State 2 local refine (EMDB - 57356) | BamGHIF Arm 2 local refine (EMDB - 57357) |
|---|---|---|---|---|---|---|---|
| **Data collection and processing** | | | | | | | |
| Magnification | 165,000 x | 165,000 x | 165,000 x | 165,000 x | 165,000 x | 165,000 x | 165,000 x |
| Voltage (kV) | 300 | 300 | 300 | 300 | 300 | 300 | 300 |
| Electron exposure (e–/Å$^2$) | 60.0 | 60.0 | 60.0 | 60.0 | 60.0 | 60.0 | 60.0 |
| Defocus range (μm) | 0.8-1.8 | 0.8-1.8 | 0.8-1.8 | 0.8-1.8 | 0.8-1.8 | 0.8-1.8 | 0.8-1.8 |
| Pixel size (Å) | 0.73 | 0.73 | 0.73 | 0.73 | 0.73 | 0.73 | 0.73 |
| Symmetry imposed | C1 | C1 | C1 | C1 | C1 | C1 | C1 |
| Initial particle images (no.) | 864,276 | 864,276 | 864,276 | 864,276 | 864,276 | 864,276 | 864,276 |
| Final particle images (no.) | 150,032 | 604,128 | 40,472 | 40,472 | 15,799 | 15,799 | 26,126 |
| Map resolution (Å) | 1.97 | 2.06 | 2.5 | 2.3 | 3.0 | 2.5 | 3.5 |
| FSC threshold | 0.143 | 0.143 | 0.143 | 0.143 | 0.143 | 0.143 | 0.143 |
| Map resolution range (Å) | 1.9-2.0 | 1.96-2.2 | 2.2-2.8 | 2.1-2.6 | 2.65-3.8 | 2.3-2.9 | 3.1-3.8 |
| **Refinement** | | | | | | | |
| Initial model used (PDB code) | - | - | - | - | - | - | - |
| Model resolution (Å) | | | | | | | |
| FSC threshold | | | | | | | |
| Model resolution range (Å) | | | | | | | |
| Map sharpening *B* factor (Å$^2$) | 42.8 | 45.9 | 31.5 | 29.0 | 27.1 | 27.2 | 49.7 |
| Model composition | | | | | | | |
| Non-hydrogen atoms | | | | | | | |
| Protein residues | - | - | - | - | - | - | - |
| Ligands | | | | | | | |
| *B* factors (Å$^2$) | - | - | - | - | - | - | - |
| Protein | - | - | - | - | - | - | - |
| Ligand | - | - | - | - | - | - | - |
| R.m.s. deviations | - | - | - | - | - | - | - |
| Bond lengths (Å) | - | - | - | - | - | - | - |
| Bond angles (°) | - | - | - | - | - | - | - |
| Validation | - | - | - | - | - | - | - |
| MolProbity score | - | - | - | - | - | - | - |
| Clashscore | - | - | - | - | - | - | - |
| Poor rotamers (%) | - | - | - | - | - | - | - |
| Ramachandran plot | - | - | - | - | - | - | - |
| Favored (%) | - | - | - | - | - | - | - |
| Allowed (%) | - | - | - | - | - | - | - |
| Disallowed (%) | - | - | - | - | - | - | - |

**Extended Data Table 2 | BamD copies used for construction of the phylogenetic tree shown in Extended Data Fig. 9d, including identifiers and mutations in the distal non-cubane Fe−S cluster (D4).** Cysteine residue annotations are based on HdrB of *Methanothermococcus thermolithotrophicus*.

| Organism | Identifier | Mutations in D4 | |
|---|---|---|---|
| *Desulfosarcina cetonica* | VTR65621 | Cys78Ser | sulfate-reducing |
| | VTR65616 | Cys78Ser | |
| | VTR63836 | Cys78Ser | |
| *Desulfotomaculum gibsoniae* | WP_006521303 | Cys78Ser | |
| | WP_006524743 | Cys78Ser | |
| | WP_015617960 | Cys78Ser | |
| | WP_006523493 | Cys78Ser | |
| | WP_006520554 | Cys78Ser | |
| | WP_006522729 | Cys78Ser | |
| *Desulfococcus multivorans* | WP_020876805 | Cys78Ser | |
| *Desulfotignum phosphitoxidans* | WP_006965651 | Cys78Ser | |
| | WP_006964228 | | |
| *Desulfotignum balticum* | WP_024335813 | Cys78Ser | |
| *Desulfomonile tiedjei* | WP_014809767 | Cys78Ser | |
| | WP_014811626 | Cys78Ser | |
| *Desulfobacula toluolica* | WP_014955904 | Cys78Ser | |
| | WP_014959477 | Cys78Ser | |
| *Desulfobacula phenolica* | WP_092238590 | Cys78Ser | |
| | WP_092236581 | Cys78Ser | |
| *Dethiosulfatarculus sandiegensis* | WP_044348230 | Cys78Ser | |
| | WP_044348930 | Cys78Ser | |
| | WP_044347634 | | |
| *Desulfocarbo indianensis* | WP_049674953 | Cys78Ser | |
| | WP_049675796 | Cys78Ser, Cys81Ser | |
| | WP_049674555 | | |
| *Desulfocucumis palustris* | WP_104372864 | Cys78Ser | |
| | WP_104372379 | Cys78Ser | |
| *delta proteobacterium NaphS2* | WP_006422201 | Cys78Ser | |
| *Geobacter daltonii (Geotalea daltonii)* | WP_012645315 | Cys78Ser | metal-reducing |
| | WP_012645331 | Cys78Ser, Cys81Ser | |
| *Geobacter bemidjiensis (Citrifermentas bemidiense)* | WP_012529877 | Cys78Ser | |
| *Geobacter metallireducens* | WP_004514581 | Cys78Ser | |
| *Geobacter pickeringii* | WP_039742563 | Cys78Ser | |
| *Geobacter pelophilus (Citrifermentans peloiphilum)* | WP_085811844 | Cys78Ser | |
| *Syntrophus aciditrophicus* | WP_011418932 | Cys78Ser | syntrophic |
| | WP_011416388 | Cys78Ser | |
| | WP_011418448 | Cys42Asp,Cys78Asp,Cys81Ala | |
| *Syntrophus gentianae* | WP_093883999 | Cys78Ser | |
| | WP_093884305 | Cys78Ser | |
| *Syntrophorhabdus aromaticivorans* | WP_028893810 | Cys78Ser | |
| | WP_051408921 | Cys78Ser | |
